# KAT6A/B inhibition synergizes with retinoic acid and enhances the efficacy of GD2-targeted immunotherapy in neuroblastoma

**DOI:** 10.1101/2025.07.01.662644

**Authors:** Nina Weichert-Leahey, Alla Berezovskaya, Mark W. Zimmerman, Francesca Alvarez-Calderon, Marlana Winschel, Silvi Salhotra, Nathaniel Mabe, Antonio Perez-Atayde, Francisca N. de L. Vitorino, Benjamin Garcia, Ulrike Gerdemann, Kimberly Stegmaier, Adam D. Durbin, Derek A. Oldridge, Brian J. Abraham, A. Thomas Look

**Author notes:** Corresponding authors: A. Thomas Look, MD, Dana-Farber Cancer Institute Department of Pediatric Oncology Mayer Bldg, Rm 630, 450 Brookline Ave, Boston, MA 02215, Brian Abraham: St. Jude Children’s Research Hospital Department of Computational Biology, Memphis, Tennessee, USA.

## Abstract

High-risk neuroblastoma accounts for about 15% of childhood cancer deaths and arises from precursors of the peripheral sympathetic nervous system. Retinoids are clinically used to inhibit growth of neuroblastoma cells through reconfiguration of the regulatory enhancer landscape. Its effects, however, are completely reversible after drug withdrawal, leading to rapid tumor cell proliferation. Here, we sought to identify epigenetic modifiers that potentiate the antiproliferative effects of retinoids in neuroblastoma. We identified PF-9363, an inhibitor of the histone H3K23 acetyltransferases KAT6A/B, as synergistically inhibiting neuroblastoma growth in combination with retinoids. PF-9363 plus retinoids induces durable growth arrest, which persists beyond retinoid withdrawal *in vitro* and *in vivo* with sustained Polycomb-mediated repression of oncogenic transcription factors MYCN, PHOX2B and GATA3. Moreover, PF-9363 plus retinoids increases GD2 expression, rendering neuroblastoma cells more sensitive to anti-GD2 immunotherapy. Overall, our studies demonstrate that KAT6A/B inhibition increases the effectiveness of retinoids and GD2-targeted immunotherapy in neuroblastoma.

## Introduction

Differentiation therapy with retinoic acid analogs has been successfully used for over two decades for the treatment of acute promyelocytic leukemia (APL) and post-consolidation therapy for pediatric high-risk neuroblastoma.^1–3^ Neuroblastomas arise from primitive neural crest cells of the developing peripheral nervous system, and are caused by mutations that i) disrupt the control of cell proliferation, often due to aberrant overexpression of *MYCN* or *MYC*; ii) block normal neuronal differentiation pathways; and iii) promote cell survival.^4–6^ Neuroblastoma is the most common extracranial solid tumor of childhood and accounts for approximately 15% of pediatric cancer-related deaths.^7^ Children with high-risk disease undergo 12 months of intensified, multimodal chemotherapy with hematopoietic stem cell rescue, surgery, and radiation therapy. Afterward, they receive six months of post-consolidation therapy, which includes isotretinoin, a retinoic acid derivative, along with antibody therapy targeting GD2, a disialoganglioside expressed on neuroectodermal tumors.^3^ This combination treatment strategy has led to improved overall survival of patients with high-risk neuroblastoma; however, relapse rates remain high, and treatment-related long-term morbidity is a serious problem.^8^

Neuroblastoma cells exhibit plasticity and are capable of interconverting between an undifferentiated neural crest-like “mesenchymal” cell state and a more differentiated sympathetic “adrenergic” cell state independent of mutation. The adrenergic cell state largely depends on a core regulatory circuitry (CRC) composed of the transcription factors (TFs) PHOX2B, GATA3, ISL1, HAND2, TBX2/3, ASCL1, SOX11, and TFAP2b, which collaborate with MYCN to drive tumor-cell growth and survival.^9–16^ Epigenetic switches from the adrenergic to the mesenchymal cell state can occur under the pressure of cytotoxic chemotherapy, or by defined genetic events.^17,18^ Therapeutic strategies to disrupt MYCN or the adrenergic TFs have been challenging due to the difficulty in attacking cell-type-specific TF proteins pharmacologically and the narrow therapeutic window for direct targeting of the affiliated transcriptional machinery.^17,19–22^

The block in neural-crest-cell differentiation that leads to uncontrolled cell proliferation in neuroblastoma can be overcome by treatment with retinoic acid, which induces neuronal differentiation, cell-cycle exit and growth arrest.^23^ After retinoic acid enters the cell nucleus, it binds to its nuclear receptor (RAR), which forms a heterodimeric complex with the retinoid X receptor (RXR). RAR-RXR receptors are bound to retinoic acid response elements (RAREs) within enhancers of target genes, and retinoic acid binding to the heterodimeric receptors results in altered gene expression. In neuroblastoma, treatment with retinoic acid downregulates members of the adrenergic CRC and MYCN by re-wiring their enhancers, and it establishes a new retino-sympathetic CRC defined by a unique set of highly expressed transcription factors (e.g., SOX4, MEIS1 and RARA), which regulate an alternative extended regulatory network of genes in neuronal progenitors that blocks proliferation and promotes differentiation.^24^ A major obstacle associated with retinoic acid therapy is that the antiproliferative effects of retinoic acid and its impact on the enhancer landscape of neuroblastoma cells are reversible once treatment is discontinued, leading to the immediate proliferation of tumor cells and disease re-occurence.^24^ Here, we sought to identify epigenetic-modifier drugs that induce durable growth arrest of neuroblastoma cells undergoing retinoic acid treatment. To this end, we performed a drug screen of 452 small-molecule inhibitors of epigenetic modifiers to identify compounds that potentiate the antiproliferative activity of retinoic acid in neuroblastoma. From this screen, we identified the KAT6A/B inhibitor PF-9363, which i) synergizes with retinoic acid to inhibit cell growth and enhances neuronal differentiation, ii) enforces sustained growth arrest *in vitro* and in murine xenograft models of neuroblastoma following retinoic acid withdrawal, as long as PF-9363 treatment is continued, and iii) enhances anti-GD2 immunotherapy. While retinoic acid treatment leads to decreased expression of the *MYCN* oncogene and of the key adrenergic transcription factors *PHOX2B* and *GATA3*, this effect is reversible. Only in combination with the KAT6A/B inhibitor PF-9363 the re-expression of these CRC members remains suppressed, thereby blocking the reactivation of cell proliferation and leading to durable growth arrest of neuroblastoma cells. In addition, retinoic acid plus PF-9363 increases cell-surface expression of GD2 on GD2^low^ neuroblastoma cells, thereby increasing tumor-cell killing by GD2-targeted antibody or chimeric antigen receptor (CAR) T cell therapy. Thus, our results demonstrate that enhancing the efficacy of anti-GD2-based immunotherapy while blocking proliferation through treatment with a KAT6A/B inhibitor plus retinoic acid is a promising novel therapeutic approach to improve outcomes for patients with high-risk neuroblastoma.

## Results

### KAT6A/B inhibition synergizes with retinoic acid to inhibit the growth of neuroblastoma

We have shown previously that the antiproliferative effects of retinoic acid treatment on *MYCN*-amplified neuroblastoma cells are due to a change in cell state through reconfiguration of their enhancer landscapes.^24^ To identify epigenetic modifiers that enhance the antiproliferative effects of retinoic acid in neuroblastoma, we conducted a drug screen using a library of 452 inhibitors of epigenetic modifiers and treated the retinoid-sensitive neuroblastoma cell line BE2C with either i) the compound alone, ii) the retinoid-derivative all-trans retinoic acid (ATRA) alone, iii) the compound in combination with ATRA, or iv) DMSO as a control (Fig. 1A). After five days of drug incubation, cell growth was determined by a CellTiter-Glo assay, and a z-score for each treatment condition was calculated to assess cell growth in comparison to the DMSO control (Fig. 1B). About one third of the compounds suppressed the growth of BE2C cells on their own, with 28 compounds showing augmentation of their antiproliferative effects due to the addition of ATRA (Fig. 1C). The compound that demonstrated the strongest potentiation of the retinoid-induced antiproliferative effects was an inhibitor of the histone demethylases (HDM) KDM6A/B, GSK-J4, which has been previously studied in combination with retinoic acid.^25^ Thus, we focused on PF-9363, which also demonstrated increased antiproliferative effects in neuroblastoma when combined with ATRA (purple bar in Fig. 1C). PF-9363 is an orally bioavailable inhibitor of KAT6A and KAT6B, which are histone acetyltransferase paralogs that acetylate H3K23 globally and H3K9 at specific gene loci.^26,27^ The clinical analog of PF-9363, PF-07248144, has recently been shown in a Phase-I trial to have significant anti-tumor activity as a single agent in patients with estrogen-receptor positive breast cancer.^28^

**Figure 1:**
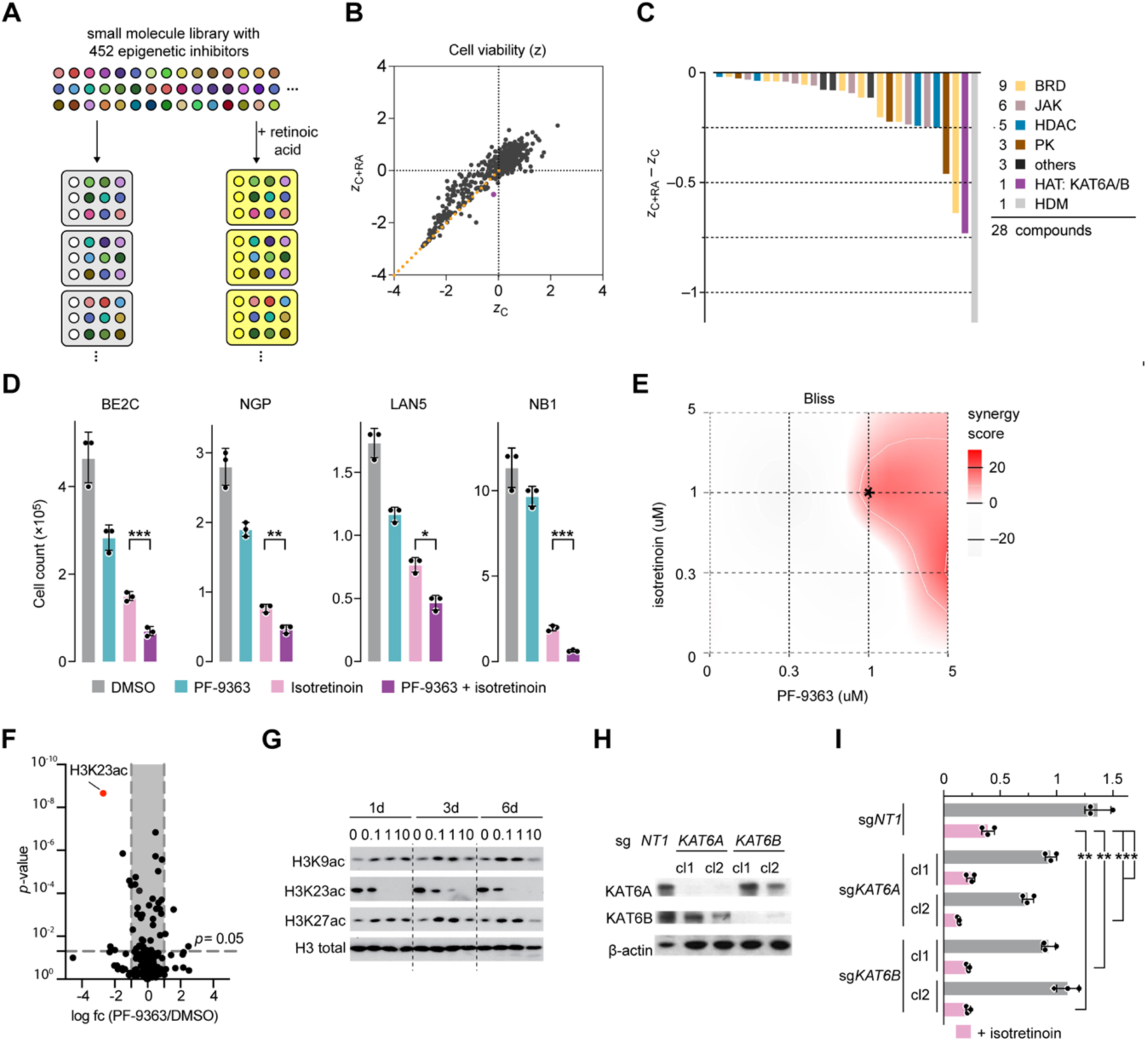
Inactivation of KAT6A and KAT6B activity synergizes with retinoic acid to suppress neuroblastoma cell proliferation. A) Schematic of a drug screen in BE2C neuroblastoma cells using small molecule inhibitors of epigenetic modifiers. BE2C cells in the white plates were treated with either DMSO control alone (clear wells in the left column) or with one of 452 compounds at a concentration of 1 μM (colored wells). In the yellow plates, BE2C cells were treated with 1 μM of the retinoic-acid-derivative ATRA alone (yellow wells in the left column) or 1 μM ATRA in combination with one of the 452 compounds at 1 μM (colored wells). After five days of treatment, viable-cell number was measured by Cell-titer Glo assay. B) Z-scores of cell viability from the drug screen in A were calculated and plotted for each compound (C) alone (x-axis) and for each compound in combination with ATRA (y-axis, C+RA). Orange dotted line shows -x = -y values, indicating no difference in growth suppression between compound alone and compound plus ATRA. Compounds plotted below the orange line exhibit more inhibition of cell growth when given in combination with ATRA than when given alone. Compound PF-9363 is marked in purple. C) Waterfall plot of the z-score for each compound in combination with ATRA minus the z-score for each compound alone (shown on the y-axis). A negative value on the y-axis indicates that the compound plus ATRA inhibits cell growth more than the compound alone, which was only the case for the 28 compounds included on the x-axis. Colored bars represent different drug targets, as outlined. Purple bar represents PF-9363, an inhibitor of the histone acetyltransferases KAT6A and KAT6B. D) Validation of the screen results using isotretinoin, the retinoic-acid-derivative that is used clinically. The neuroblastoma cell lines BE2C, NGP, LAN5 and NB1 were each treated with DMSO, 1 μM PF-9363 alone, 1 μM isotretinoin alone, or both compounds at 1 μM each for eight days. Shown are cell counts from three independent experiments. Significance for the combination treatment compared with other treatment groups was determined by an unpaired t-test. \**p*< 0.05, ** *p*< 0.01, \*\*\**p*< 0.001. E) Synergy map created using the publicly available software SynergyFinder+ and the synergy-scoring method called BLISS, which revealed synergistic effects (red zone) for the combination of isotretinoin plus PF-9363 after seven days of treatment in BE2C cells. Asterisk indicates BLISS synergy score >15 for this drug combination when each compound is given at 1 μM. F) Volcano plot of log fold changes of histone proteins measured by mass spectrometry extracted from BE2C cells treated with PF-9363 1 μM for 14 days versus DMSO control treated cells. G) Western blot analysis using antibodies to the indicated histones that were extracted from neuroblastoma cell line KELLY after treatment with the indicated doses of PF-9363 for 1, 3 and 6 days. Total H3 served as a loading control. H) Western blot assay measuring KAT6A and KAT6B protein levels in BE2C cells that had been stably transduced with Cas9 and a sgRNA either targeting *KAT6A* (*sgKAT6A*) or *KAT6B* (*sgKAT6B*). BE2C cells with CRISPR-Cas9 guided knockout of a non-targeting sequence (NT1)^24^ were used as a control cell line. Beta actin was measured as a loading control. I) Absolute cell counts of NT1 (grey bar), *sgKAT6A* or *sgKAT6B* BE2C cells (light grey bars for clone 1 and clone 2). Cells were treated with either DMSO or 1μM isotretinoin (pink bars) for seven days. Significance between groups treated with isotretinoin was determined by an unpaired t-test; \**p*< 0.05, \*\**p*< 0.01.

To validate the results of the drug screen, we tested the effects of PF-9363 alone and in combination with the retinoic acid-derivative isotretinoin, which is used clinically in neuroblastoma, using several retinoid-sensitive neuroblastoma cell lines. We treated BE2C, NGP, LAN5 and NB-1 cells with either DMSO control, isotretinoin alone, PF-9363 alone, or the combination of PF-9363 plus isotretinoin and measured cell growth (Fig. 1D). As predicted from our screen, treatment with the combination of isotretinoin plus PF-9363 suppressed the growth of neuroblastoma cells significantly more than treatment with either isotretinoin or PF-9363 alone (Fig. 1D). Reduced tumor cell number among all four treatment groups was due to proliferative arrest and was not caused by apoptosis (Supplementary Fig. S1). To test whether the antiproliferative effects of each drug were synergistic or additive, we conducted excess over BLISS synergy analysis.^29^ By testing the effects of several concentrations of PF-9363, alone and in combination with isotretinoin, on BE2C cell numbers, we found that after seven days of treatment, the drug combination was synergistic over a wide range of concentrations (highlighted in red), including 1 μM for each compound (Fig. 1E, asterisk), with an excess-over-BLISS synergy score of >15.

To test whether treatment with 1μM of PF-9363 exclusively inhibits the acetylation of H3K23 in neuroblastoma through specific inhibition of KAT6A/B, and that no other nucleosome-modifying enzymes are affected at this dose, we performed mass spectrometry to detect changes of all histone modifications following PF-9363 treatment. After 14 days of treatment with 1μM of PF-9363, H3K23ac was the most significantly downregulated histone modification compared to DMSO treatment, and no other single-residue mark was altered (Fig. 1F). To confirm these mass spectrometry findings, we extracted histones from neuroblastoma cells treated with several concentrations of PF-9363 for 14 days and performed western blotting with an antibody specific for H3K23 acetylation (ac) (Fig. 1G). We also analyzed the amount of H3K27ac, which is catalyzed by KAT3A/B (EP300/CBP), and H3K9ac, which has been implied in previous studies to be facilitated by KAT6A/B at specific gene loci.^30^ Inhibition of global H3K23ac by PF-9363 was observed at 1μM, but we did not observe reduced levels of H3K9ac or H3K27ac at 1μM (Fig.1G), indicating that at this concentration, PF-9363 specifically blocks KAT6A/B-mediated acetylation of H3K23 without affecting the other enzymes.^31,32^

To compare the effects of pharmacological inhibition of KAT6A and B using PF-9363 with the effects of genetic knockout of either enzyme, we disrupted the endogenous *KAT6A* and *KAT6B* genes using a CRISPR-Cas9 system (Fig.1H). Next, the growth of BE2C cells that had been stably transduced with Cas9 and a sgRNA targeting either i) *KAT6A (*sg*KAT6A)* or ii) *KAT6B (*sg*KAT6B)* was measured after treatment for seven days with isotretinoin alone (pink bars) or DMSO (grey bars) as a control (Fig.1I).^24^ We found that sg*KAT6A* and sg*KAT6B* BE2C cells showed reduced growth compared to the control BE2C cells given a non-targeting guide (*NT1*), indicating that each of these gene paralogs contributes to optimal cell growth. As observed for the pharmacologic inhibition of KAT6A/B activity, cell growth inhibition of sg*KAT6A* and sg*KAT6B* BE2C cells was more pronounced when they were treated with isotretinoin (Fig.1I).

### KAT6A and KAT6B activity is necessary for neuroblastoma cellular identity

The roles of KAT6A/B-facilitated acetylation of H3K23 in regulation of chromatin structure and gene expression are emerging and the subjects of ongoing investigations.^33–35^ To dissect the roles of KAT6A/B in chromatin accessibility and transcriptional regulation in neuroblastoma, we assessed chromatin binding sites of each enzyme and correlated their binding profiles with profiles of H3K27ac and ATAC-seq. Here, we found that most genome-wide KAT6A and KAT6B binding was localized at promoters rather than at regulatory enhancers (Fig. 2A). At those promoters, KAT6A and KAT6B generate H3K23 acetylation, which occurs preferentially downstream of the transcription start site (TSS; Fig. 2B,C); both marks were locally enriched immediately adjacent to ATAC regions of transpose-accessible DNA (Fig. 2B, C). The asymmetric enrichment for H3K23Ac modifications and open chromatin downstream of TSS may result from promoter-proximal pausing of RNA polymerase II (PolII), which often occurs after initiating transcription, typically around 20-60 bases downstream of the transcription start site.^36^. Loss of either KAT6A or KAT6B in the sg*KAT6A/B* BE2C cell lines led to decreased H3K23ac at these promoter regions (Fig. 2B). Thus, both KAT6A and KAT6B preferentially act at active promoters downstream of the TSS, supporting a role of H3K23 acetylation in chromatin accessibility at promoters and of KAT6A and KAT6B during transcriptional regulation in neuroblastoma.

**Figure 2:**
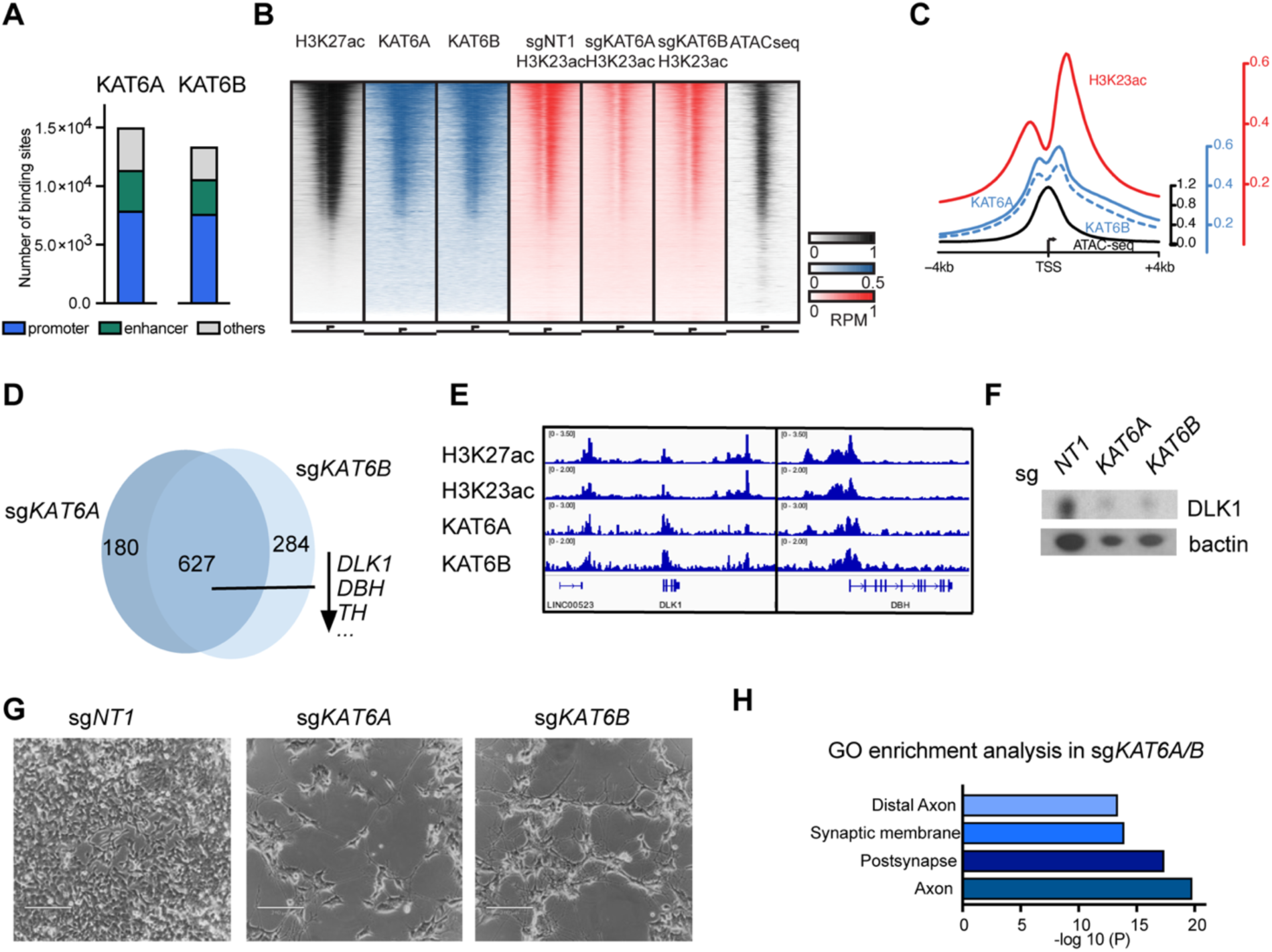
Mechanistic roles of KAT6A/B in genome regulation. A) Genome-wide distribution of significant KAT6A and KAT6B CUT&RUN peaks. B) Reads-per-million (RPM) normalized promoter coverage heatmap for relevant chromatin marks. 2kb regions centered on all transcription start sites were split equally into 50 bins and ranked by H3K27ac signal. A strong correlation among all marks at the subset of active promoters is apparent. Loss of H3K23ac in BE2C cells with sg*KAT6A* and sg*KAT6B* shown with sg*NT1* used as a control. C) Reads-per-million normalized average coverage at all promoters (same regions and parameters as B). Each mark is separately scaled using distinct Y axis on right for visibility. Histone modifications appear adjacent the transcription start site and its associated nucleosome-free region. D) Venn diagram indicating numbers of genes differentially expressed in both BE2C sg*KAT6A* and sg*KAT6B* with log fc <-0.5. E) Gene tracks at *DLK1* and *DBH* gene loci with H3K27ac, H3K23ac, KAT6A and KAT6B ChIP-Seq/CUT&RUN. F) Western blot using protein extractions from BE2C sg*NT1*, sg*KAT6A*, sg*KAT6B* using antibody against DLK1 and β-actin as loading control. G) Morphology of BE2C sg*NT1*, sg*KAT6A*, sg*KAT6B* cells assessed by dark-field microscopy. H) Gene Set Enrichment Analysis of genes upregulated in both BE2C sg*KAT6A* and sg*KAT6B* cells (log2 fc>0.5) versus control BE2C sg*NT1* cells. Gene Ontology (GO) sets shown here are among the top 10 sets in the enrichment analysis.

To further explore the cellular mechanisms that underpin the antiproliferative effects of KAT6A and KAT6B inhibition and its synergism with retinoic acid in neuroblastoma (Fig.1), we performed bulk RNA-seq analysis from sg*KAT6A* and sg*KAT6B* BE2C cells. Here, we specifically examined the commonalities in the transcriptional programs regulated by KAT6A and KAT6B and found a large overlap of significantly downregulated genes in both sg*KAT6A* and sg*KAT6B* BE2C cells (Fig. 2D). Among the genes that showed differential expression in both the sg*KAT6A* and sg*KAT6B* BE2C cells, serval key neuroblastoma identity genes were downregulated, including the *dopamine beta-hydroxylase* (*DBH*), *tyrosine hydroxylase (TH)* and *Delta-like canonical notch ligand 1* (*DLK1*) (Fig. 2D). As these genes were co-regulated in both sg*KAT6A* and sg*KAT6B* cell lines, we next sought to understand if these gene expression changes were a direct consequence of chromatin occupancy of these enzymes. Chromatin pulldown data revealed direct binding of both KAT6A and KAT6B at the promoters of *DBH* and *DLK1*, identifying them as direct targets of both KAT6A and KAT6B in neuroblastoma (Fig. 2E). Given the established role of DLK1 in maintaining a more immature, undifferentiated state of neuroblastoma,^37^ the reduced expression of DLK1 upon KAT6A/B loss (Fig. 2F) likely contributes to the observed i) reduced growth (Fig. 1I), ii) promotion of morphological differentiation (Fig. 2G) and iii) transcriptional enrichment for gene signatures characteristic of neuronal differentiation observed in sg*KAT6A* and sg*KAT6B* BE2C cells compared to sg*NT1* Be2C cells (Fig. 2H). Thus, these results emphasize that both KAT6A and KAT6B entertain a central role in maintaining neuroblastoma proliferation and suppressing neuronal differentiation through the transcriptional regulation of key neuroblastoma genes with distinct signatures.

### Retinoid-induced neuroblastoma differentiation is augmented by KAT6A/B inhibition

To further investigate mechanisms of synergy between retinoic acid and PF-9363 treatment and to account for heterogeneity in cellular responses to drug treatments, we performed single-cell RNA sequencing (scRNA-seq). Treatments groups included i) DMSO, ii) isotretinoin, iii) PF-9363, or iv) the combination of PF-9363 and isotretinoin, all treated for 14 days prior to scRNA-seq. Uniform Manifold Approximation and Projection (UMAP) analysis revealed two major groups primarily containing cells treated with isotretinoin (pink bracket) versus no isotretinoin (blue bracket), revealing that retinoid treatment transforms adrenergic cells into new, non-overlapping cell states (Fig. 3A). Strikingly, cells treated with either isotretinoin alone or in combination with PF-9363 resided together in one major group, independent of PF-9363 treatment (Fig. 3B, Supplementary Fig. S2A). Next, we conducted unsupervised clustering analysis using all treatment groups and identified eight distinct cell clusters, including clusters 1,5 and 6 enriched in the ‘no isotretinoin’ group and clusters 2-4, 7 and 8 enriched in the ‘isotretinoin-treated’ group (Fig. 3C, D). This separation of clusters underscores the dominant influence of isotretinoin on the transcriptional landscape, regardless of PF-9363 co-treatment. However, the addition of PF-9363 treatment to isotretinoin led to loss of cells in cluster 2, where almost half of all isotretinoin-treated cells reside, and increased the size of clusters 3 and 4, which account for two thirds of all cells treated with PF-9363 plus isotretinoin (Fig. 3E, Supplementary Fig. S2B).

**Figure 3:**
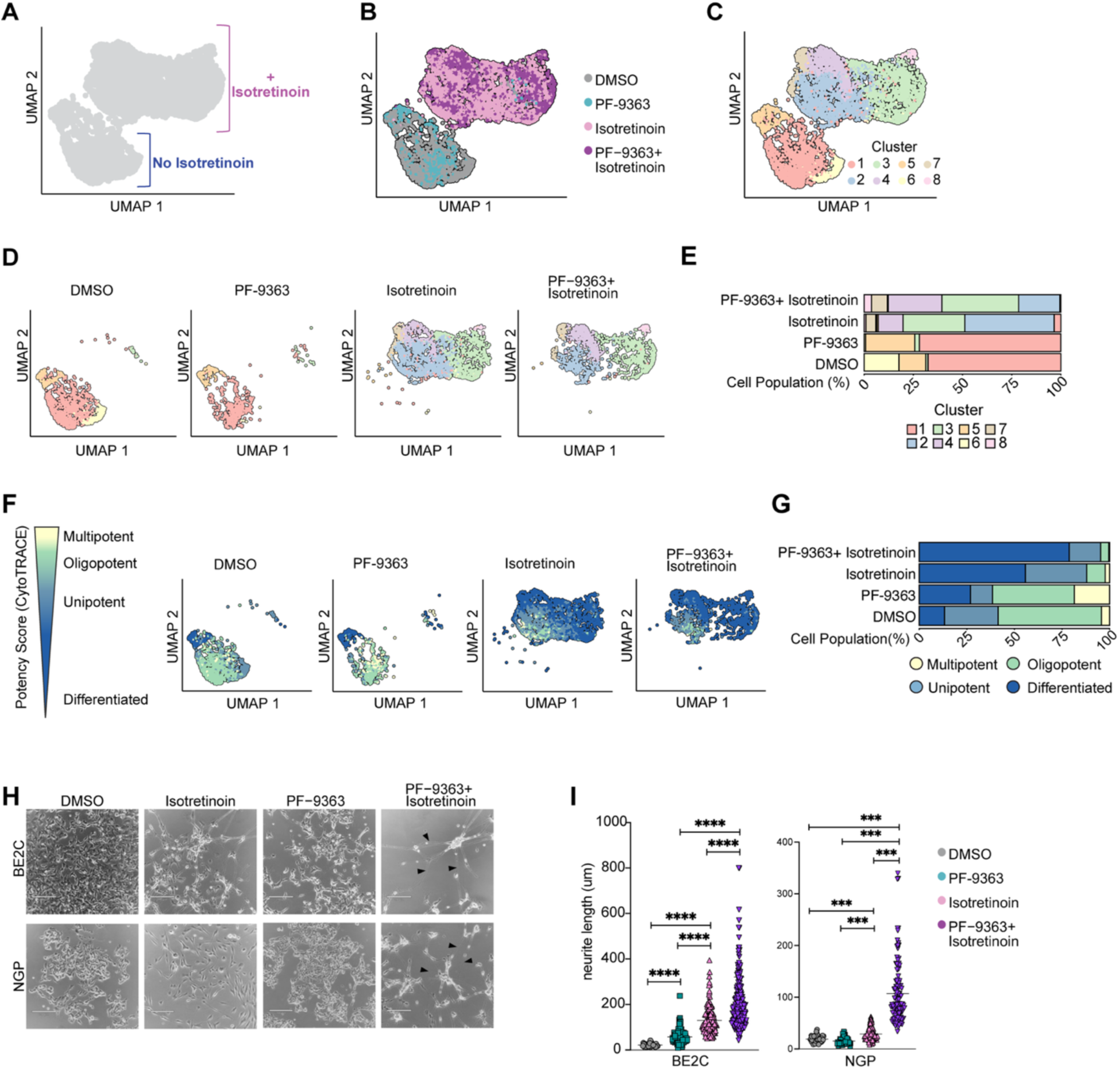
The combination of KAT6A/B inhibition and retinoic acid enhances differentiation in neuroblastoma cells. Neuroblastoma BE2C cells were treated with either DMSO, PF-9363 alone, isotretinoin alone, or isotretinoin plus PF-9363. Each compound was administered at a concentration of 1 μM. A) Single-cell RNA-seq was performed and UMAP analysis conducted across all treatment groups merged together. B) UMAP analysis from BE2C cells highlighting all four treatment conditions. C) Cluster analysis was performed including cells from all four treatment groups, which identified 8 different clusters. D) Cluster distribution in each treatment condition on the UMAP. E) Bar chart of cluster distribution within each treatment group expressed as % cell population. F) CytoTRACE 2 potency analysis was performed in single cells from all four treatment groups, here shown separately for each treatment condition. G) Bar chart with distribution of cell potency in % contributing to each treatment group. H) Morphology of BE2C and NGP cells after 14 days of treatment was assessed by dark-field microscopy. Scale bar = 210 μm. Black arrow heads point to neurites. I) Neurite length (in μm) was measured in BE2C and NGP cells after 14 days of treatment. Significance was determined using the t-test. *** *p< 0.001,* **** *p< 0.0001*.

To evaluate differences in cellular differentiation across treatment groups, we used CytoTRACE2, which classifies cells over a range of cellular potencies including ‘multipotent’, ‘oligopotent’, ‘unipotent’, and ‘differentiated’.^38^ In the DMSO-treated control experiment, most cells derived from cluster 1 (Fig. 3E), which was the least differentiated of all clusters and primarily classified as ‘oligopotent’ (Fig. 3F, Supplemental Fig. S2C), indicating a relatively undifferentiated state consistent with neuroblasts. A smaller population of cells fell into cluster 6 (Fig. 3D,E), which was primarily ‘unipotent’ (Fig. 3F, Supplemental Fig. S2C), reflecting the baseline heterogeneity of untreated neuroblastoma cells. In cells treated with PF-9363 alone, cluster 1 was the dominant cluster, including ‘oligopotent’ cells and a new population of ‘multi-potent’ cells. However, cells from cluster 6 largely disappeared upon PF-9363 treatment (Fig. 3D,E), while cluster 5 increased with more ‘differentiated’ cells due to PF-9363 treatment (Fig. 3F, G, Supplemental Fig. S2B, C). These findings suggest that PF-9363 may act as a differentiation agent, promoting differentiation in a subset of cells that have already begun to lose some developmental potency, including those cells that move from cluster 6 to cluster 5.

Despite the major transformation of cell state through isotretinoin treatment alone (Fig. 3A) and an overrepresentation of isotretinoin treated cells in cluster 2 (Fig. 3D, E, Supplemental Fig. S2B), only ∼60% of the treated cells adopted a ‘differentiated’ phenotype, while the remaining cells remained of higher potency, including oligo- and multipotency (Fig. 3F, G). By contrast, upon combination treatment with isotretinoin plus PF-9363, representation of cluster 2 markedly decreased, and the amount of differentiated cells increased to ∼ 80%, primarily affecting cells in clusters 3, 4, 7, and 8 (Fig. 3C-E, Supplemental Fig. S2C). Importantly, the combination treatment is much more effective in decreasing the number of oligopotent and multipotent cells than isotretinoin treatment alone, which presumably represent the cells that cause regrowth of dividing neuroblasts upon isotretinoin withdrawal.

The morphological changes of neuroblastoma cells undergoing treatment with retinoic acid alone or in combination with PF-9363 were consistent with the transcriptional changes we observed on the single-cell level. PF-9363 treatment alone induced subtle morphological changes over 14 days of treatment, in which neuroblastoma cells formed very short neurites (Fig. 3H, I). Isotretinoin treatment induced morphological differentiation of neuroblastoma cells, as previously reported,^23^ with the formation of short neurites in both BE2C and NGP cells (Fig. 3H,I). By contrast, the combination treatment with isotretinoin and PF-9363 induced significant changes in the phenotype of both BE2C and NGP cells (Fig. 3H), with the formation of significantly longer neurites (Fig. 3I).

These results highlight the limitations of isotretinoin monotherapy in neuroblastoma. Moreover, our data suggest a model in which isotretinoin relieves an early block in differentiation, priming cells for further maturation. However, isotretinoin alone is insufficient to drive terminal differentiation for all cells. Conversely, co-treatment with the KAT6A/B inhibitor PF-9363 largely overcomes these limitations, markedly enhancing neuronal differentiation both morphologically and at the transcriptional level compared to isotretinoin alone. PF-9363 appears to act on cells already primed for differentiation, pushing them toward a more fully differentiated state when given in combination with isotretinoin. These results provide insight into the mechanism of synergy, with isotretinoin initiating and PF-9363 accentuating differentiation, resulting in differentiation of a larger proportion of treated neuroblasts.

### KAT6A/B inhibition prolongs the antiproliferative effects of retinoic acid in neuroblastoma

Next, we assessed whether the administration of a KAT6A/B inhibitor in combination with isotretinoin could not only augment cell growth arrest and neuronal differentiation of malignant neuroblasts, but also potentially prolong the growth suppression of neuroblastoma cells following the withdrawal of isotretinoin. Since isotretinoin, a component of the standard post-consolidation treatment for patients with high-risk neuroblastoma in North America, is limited to a 14-day treatment length per cycle to minimize toxicity, we aimed to replicate this schedule in our *in vitro* experiments. In these experiments, we monitored the growth of cells treated with either i) isotretinoin alone for 14 days, ii) PF-9363 alone for 28 days, iii) isotretinoin plus PF-9363 for 14 days, followed by withdrawal of both drugs, and iv) isotretinoin plus PF-9363 for 14 days, followed be removal of isotretinoin and continuation of PF-9363 for the entire 28 days (Fig. 4A). PF-9363 was administered for the entire 28-day period, based on the tolerability profile of the clinical lead compound PF-07248144, for which daily administration was reported to be safe in a recent clinical trial.^28^ Neuroblastoma cells (BE2C and NGP) treated with PF-9363 only for 28 days showed exponential growth, although at a slower rate compared to cells treated with DMSO (group I versus II in Fig. 4A). In neuroblastoma cells treated with isotretinoin alone for 14 days, followed by DMSO for 14 days, cell growth was dramatically suppressed by isotretinoin for the initial 14-day period (pink line, Fig. 4A, group III) and cells appeared morphologically differentiated (Supplementary Fig. S3). However, after isotretinoin was removed, rapid resumption of growth occurred and the cells reverted to small round proliferating neuroblastoma cells (Supplementary Fig. S3), indicating that the antiproliferative effects of isotretinoin alone are rapidly reversible upon drug withdrawal. During treatment with isotretinoin in combination with PF-9363, cell growth was significantly more suppressed (purple line, Fig. 4A) than by isotretinoin alone. However, a resumption of cell growth was observed after both drugs were removed (Fig. 4A, group IV). By contrast, when only isotretinoin was discontinued, but PF-9363 was continued for an additional 14 days, neuroblastoma cell growth was continuously suppressed (Fig. 4A, group V) and cell morphology continued to be differentiated (Supplementary Fig. S3).

**Figure 4:**
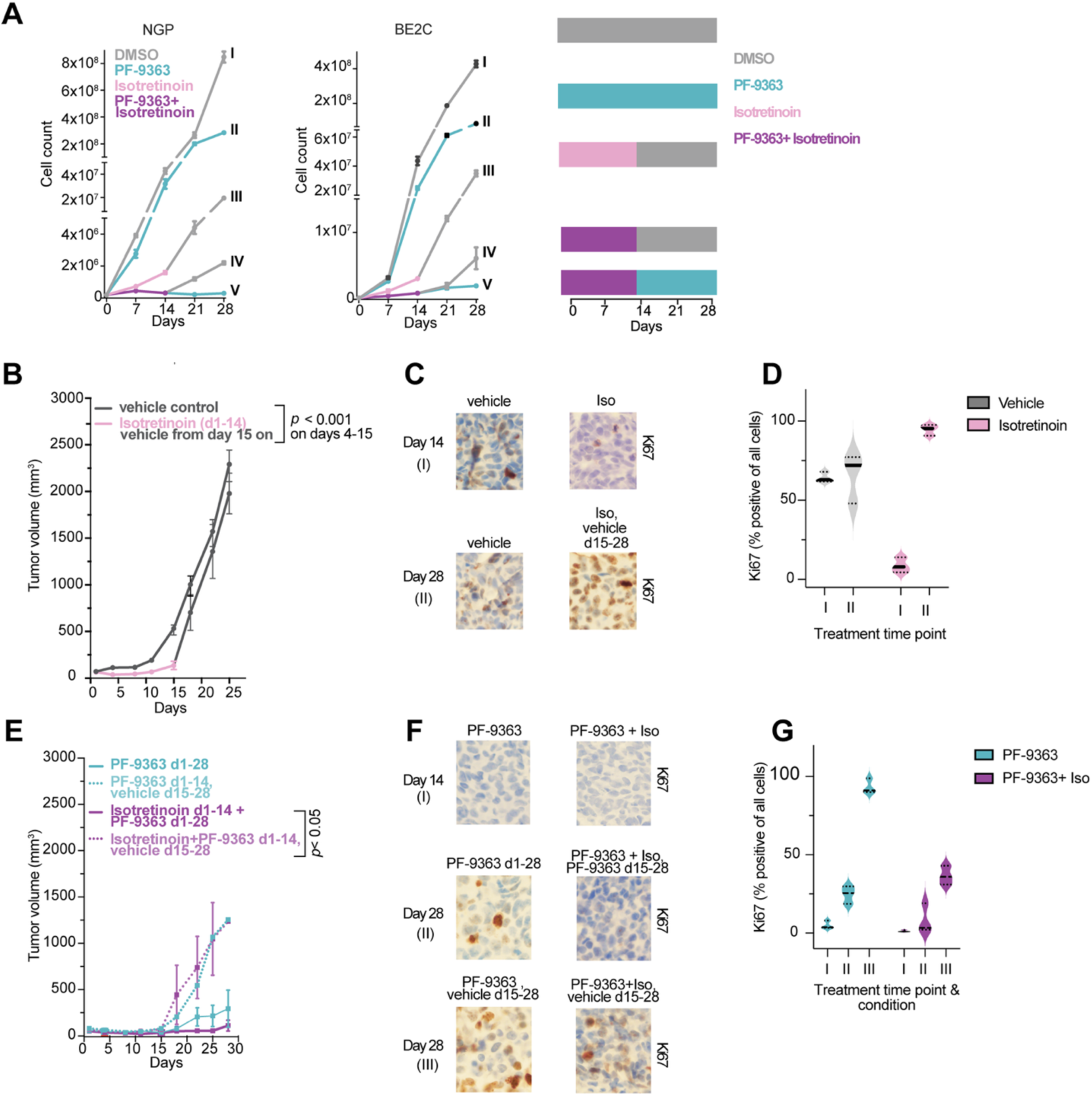
KAT6A/B inhibition prolongs retinoid-induced growth suppression in neuroblastoma. A) Representative growth curve for BE2C cells treated with either I) DMSO control (continuous grey line) for 28 days, II) PF-9363 alone (continuous green line) for 28 days, isotretinoin for 14 days (pink line fragment) followed by a 14-day drug-washout period with DMSO (grey line fragment), IV) PF-9363 plus isotretinoin for 14 days (purple line fragment) followed by a 14-day drug-washout period with DMSO (grey line fragment), or V) PF-9363 plus isotretinoin for 14 days (purple line fragment) followed by a 14-day isotretinoin-washout period during which PF-9363 was continued (green line fragment). Each compound was administered at a concentration of 1 μM. B) BE2C neuroblastoma cell xenografts were established in NSG mice. Mice were then treated with vehicle control (*n* = 9) for 28 days (continuous grey line), or isotretinoin for 14 days (pink line fragment) followed by vehicle control until day 28 (grey line fragment, *n = 9*). Tumor volume was significantly different between vehicle and isotretinoin treatment groups on days 4 through 15 (*p* < 0.001 as determined by Mann-Whitney test). Data are shown as mean values ± SEM. C) Immunohistochemistry (IHC) for Ki67 was performed on FFPE tissue sections from BE2C xenograft tumors at the indicated time points and treatments corresponding to the experiment described in panel A. D) Quantification of Ki67 positive cells in C. E) BE2C-xenografted mice were treated with either PF-9363 for 28 days (green line, *n* = 5), PF-9363 for 14 days followed by vehicle control for 14 days (dotted green line, *n* = 4), isotretinoin plus PF-9363 for 14 days followed by vehicle control for 14 days (dotted purple line, *n* = 4), or isotretinoin plus PF-9363 for 14 days followed by PF-9363 only for 14 days (purple line, *n* = 5). As determined by a Mann-Whitney test, tumor volume was significantly different (*p* < 0.05) on days 18, 22 and 25 when comparing treatment groups isotretinoin plus PF-9363 d1-14 followed by vehicle d15-28 (dotted purple curve) and isotretinoin plus PF-9363 d1-14 followed by PF-9363 d15-28 (purple curve). Data are shown as mean values ± SEM. F) IHC for Ki67 was performed on FFPE tissue sections from BE2C xenografts at the indicated time points and treatments corresponding to the experiment described in panel C. G) Quantification of Ki67 positive cells in F.

To study the efficacy of isotretinoin treatment on neuroblastoma cell growth *in vivo*, we established subcutaneous xenografts of BE2C cells in the flanks of NSG mice. Four days after tumor implantation, mice were treated with either vehicle control or isotretinoin at 50 mg/kg once daily by oral gavage for 14 days. Isotretinoin treatment inhibited tumor growth over the 14-day period, with significantly smaller tumor volumes in treated mice compared to those receiving the vehicle alone, starting at day 4 of treatment and continuing through to the end of treatment on day 14 (Fig. 4B). Interestingly, engrafted tumor cells did not demonstrate morphologic evidence of neuronal differentiation when treated with isotretinoin in the xenograft BE2C model, which is different from our *in vitro* results. After 14 days of treatment, isotretinoin was stopped, and the treatment was changed to vehicle control. Consistent with our *in vitro* results (Fig.4A), tumors began to grow immediately after isotretinoin treatment was discontinued and tumor volumes increased rapidly (Fig.4B), indicating the transient nature of isotretinoin-induced growth suppression *in vivo* and providing a plausible mechanism for the lack of long-term benefits from isotretinoin treatment for patients with high-risk neuroblastoma.^3^

To explore the underlying mechanism of suppressed tumor growth in mice treated with isotretinoin, tumors were excised, formalin-fixed and paraffin-embedded (FFPE), and sectioned for immunohistochemistry (IHC) with an antibody against Ki67. Tumors from the vehicle control group showed a high fraction of nuclei stained for Ki67 at both days 14 and 25, indicating a high fraction of cells in S phase of the cell cycle, whereas sections of tumors from mice treated with isotretinoin alone for 14 days exhibited a very low fraction of nuclei staining positive for Ki67 at day 14 (Fig. 4C, D). By contrast, tumors from animals that were treated with isotretinoin alone for 14 days followed by vehicle for 14 days showed a high fraction of Ki67-positive nuclei at day 25 (Fig.4C, D), indicating a complete reversal of the isotretinoin-induced proliferative arrest almost two weeks after treatment cessation.

To test the efficacy of PF-9363 as a single agent for the treatment of neuroblastoma *in vivo*, BE2C-xenografted NSG mice were treated with 5 mg/kg PF-9363 daily by oral gavage for either 14 or 28 days. PF-9363 treatment showed significant anti-tumor activity that was similar to isotretinoin (Fig. 4E), which was unexpected based on its mild antiproliferative effects on neuroblastoma cells growing *in vitro* (Fig. 4A). This difference is apparently due to the fact that the PF-9363 concentration of 1 µM *in vitro* inhibits only KAT6A/B, while PF-3963 administered at 5 mg/kg daily to mice *in vivo* inhibits both KAT6A/B and KAT7.^31,32^ When PF-9363 was discontinued after 14 days of treatment, tumor cells grew rapidly for the remaining 14 days of the experiment (dotted green curve in Fig. 4E), and even with continued treatment until day 28 (green curve in Fig. 4E), tumor cells began to slowly grow. Ultimately, tumor volumes were not significantly different between these two treatment groups.

To assess the S-phase fraction in this treatment group with PF-9363 alone, tumor sections were stained for Ki67 at days 14 and 28. Tumor cell nuclei were Ki67-negative at day 14; however, at day 28, the fraction of Ki67+ tumor-cell nuclei was modestly increased in tumors that had been treated continuously with PF-9363 (Fig. 4F, G) and much higher in tumors for which PF-9363 treatment was discontinued after day 14 (Fig. 4F, G), which is consistent with the observed increase in tumor volume (Fig. 4E). Thus, PF-9363 treatment alone at 5 mg/kg once daily to mice initially mediates promising growth suppression of xenografted neuroblastoma cells *in vivo*; however, its antiproliferative effect as a single agent is progressively lost after 14 days of treatment.

To test whether the combination treatment with isotretinoin plus PF-9363 induces sustained growth suppression of neuroblastoma cells *in vivo*, we treated tumor-bearing NSG mice with the combination of isotretinoin and PF-9363 for 14 days and observed suppression of tumor growth similar to that of either drug alone (Fig. 4B, 4E). When both compounds were discontinued on day 14, tumors began to grow rapidly (Fig. 4E, purple dotted curve). However, in contrast to single-agent treatment, we observed sustained growth suppression for the 28-day treatment period when only isotretinoin was stopped after 14 days and PF-9363 was continued (Fig. 4E, purple curve), and tumor volumes were significantly different between these two treatment groups.

An analysis of S-phase fractions in treated tumor cells showed that mice receiving both drugs for 14 days lacked Ki67-positive nuclei on day 14 (Fig. 4F, G), indicating the absence of tumor-cell proliferation. Tumors from mice, in which both isotretinoin and PF-9363 were withdrawn at day 14, stained strongly positive for Ki67 on day 28 (Fig. 4F, G), consistent with rapid tumor growth after stopping both compounds. However, tumors from mice treated with both drugs for 14 days and then with PF-9363 only during days 15-28 showed very low levels of Ki67 staining (Fig. 4F, G), which is consistent with the profound suppression of cell growth (Fig. 4E). Taken together, tumor re-growth after isotretinoin withdrawal was shown to be due to tumor-cell proliferation marked by Ki67 positivity, and both neuroblastoma cell growth and Ki67 nuclear positivity was prevented by KAT6A/B inhibition during days 15 to 28.

Thus, in our pre-clinical xenograft model of neuroblastoma, the novel combination treatment with isotretinoin plus the KAT6A/B inhibitor PF-9363 is well tolerated, without excessive weight loss (Supplementary Fig. S4), and induces durable growth arrest of the neuroblastoma cells, which persists after isotretinoin withdrawal, as long as daily PF-9363 is continued.

### Sustained down-regulation of adrenergic transcription factors by treatment with retinoic acid plus KAT6A/B inhibition

Retinoic acid treatment alone rewires the enhancer landscape in neuroblastoma, temporarily changing the transcriptional circuitry from the proliferative adrenergic CRC to the non-proliferative retino-sympathetic CRC.^24^ To investigate the epigenetic changes and the enhancer landscape underlying the sustained growth suppression of neuroblastoma cells by treatment with isotretinoin and the KAT6A/B inhibitor PF-9363, we performed chromatin pulldown using an antibody against H3K27 trimethylation, a histone modification associated with gene silencing and repressed expression, in BE2C cells from each treatment group. In the analysis of H3K27me3 deposition, only two adrenergic CRC members were among the gene loci that showed the greatest increase in H3K27me3 (log fold change>1) upon isotretinoin treatment alone or in combination with PF-9363: *PHOX2B* and *GATA3* (Fig. 5A). Importantly, the regulatory elements of both *PHOX2B* and *GATA3* showed dramatically increased amounts of H3K27me3 deposition upon combination treatment compared to single-agent treatment (Fig.5B, C).

**Figure 5:**
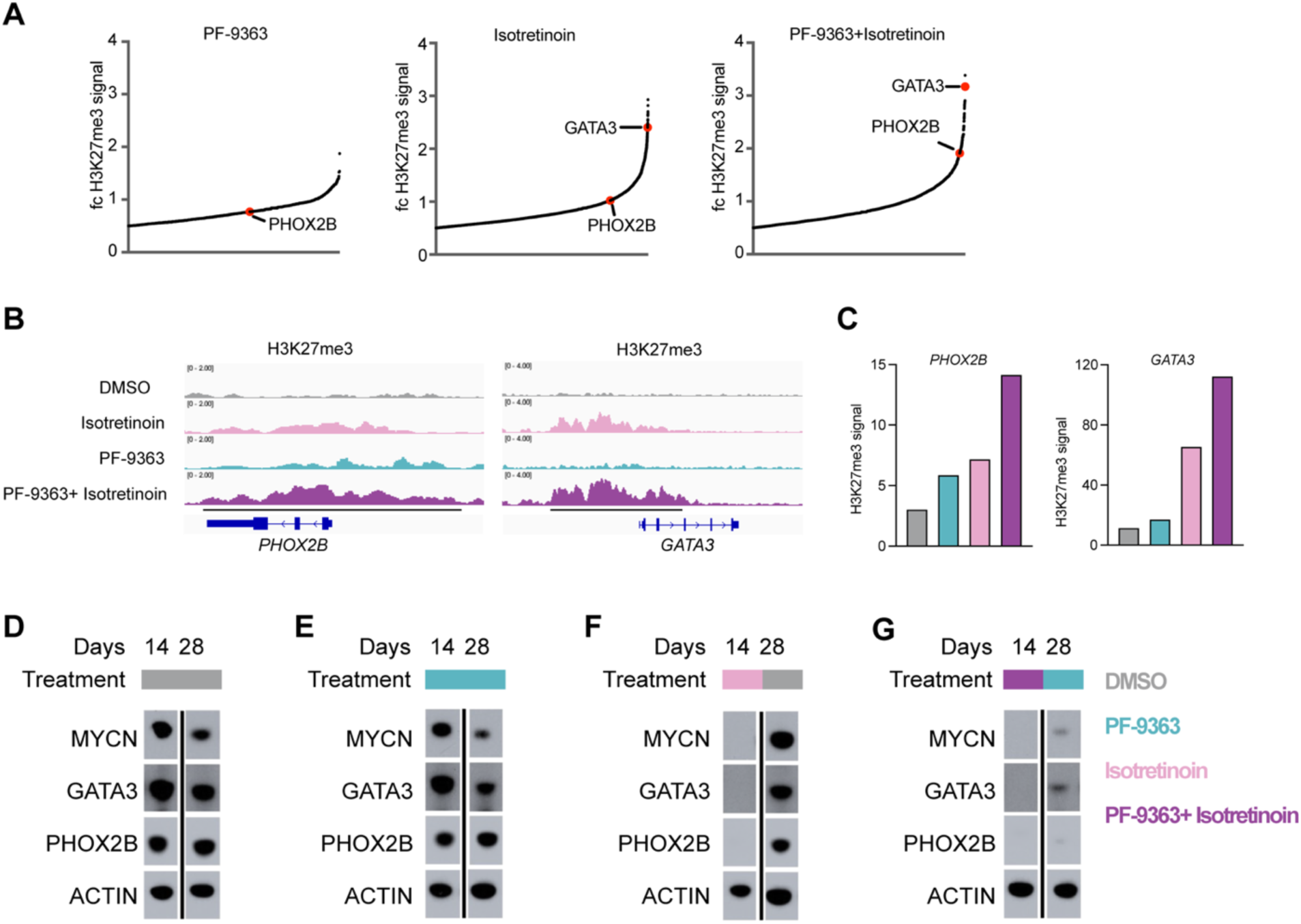
KAT6A/B inhibition prolongs retinoid-induced suppression of gene expression of adrenergic CRC members. A) Scatter plot of log2 fold-changes in H3K27me3 CUT&RUN signal at significant islands with lfc > 0.5 versus their fold-change rank. Experiments obtained from BE2C cells treated with PF-9363, isotretinoin or isotretinoin plus PF-9363 for 14 days at a concentration of 1 μM compared to control DMSO-treated BE2C cells. Significant islands associated with *PHOX2B* and *GATA3* are noted. B) Gene tracks at *PHOX2B* and *GATA3* gene loci from CUT&RUN experiments with antibody against H3K27me3 in BE2C cells treated with DMSO, isotretinoin, PF-9363 or isotretinoin plus PF-9363 for 14 days. C) Bar chart of quantified H3K27me3 signal in the regulatory elements (indicated as black line in panel B) of the *PHOX2B* and *GATA3* gene loci. D-G) Western-blot assay for transcription factors GATA3, PHOX2B and MYCN in BE2C cells treated with either DMSO as a control for 28 days (D), PF-9363 for 28 days (E), isotretinoin for 14 days followed by DMSO for 14 days (F), or isotretinoin plus PF-9363 for 14 days followed by PF-9363 alone for 14 days (G). Each compound was administered at a concentration of 1 μM. β actin was included as a loading control. Western blots were confirmed in two independent experiments. Cropped images are shown here. Complete Western blots are in Supplementary Figure 6.

To determine whether the combination treatment–mediated increase in H3K27me3 at the regulatory elements of *PHOX2B* and *GATA3* would lead to reduced expression of these transcription factors and persist after isotretinoin removal, we assessed the protein levels of PHOX2B, GATA3, and MYCN at serial time points. When neuroblastoma cells were treated with PF-9363 alone for 14 and 28 days, we observed sustained high expression levels of MYCN, PHOX2B and GATA3, similar to the DMSO control cells (Fig. 5D, E), which is consistent with the growth rate of neuroblastoma cells treated with PF-9363 alone (group II in Fig. 4A). Isotretinoin treatment for 14 days led to dramatical downregulation of the expression levels of MYCN, PHOX2B and GATA3 (Fig. 5F), which is consistent with the growth arrest observed in these cells (group III in Fig. 4A). However, 14 days after isotretinoin was removed, MYCN, PHOX2B and GATA3 were again expressed at high levels (Fig. 5F), similar to control cells (Fig. 5D), mediating the rapid return of cell proliferation with this treatment (Fig. 4A). In contrast, in BE2C cells that were treated initially with isotretinoin plus PF-9363 followed by withdrawal of isotretinoin but continuous exposure to PF-9363 for the entire 28 days, protein levels of MYCN, PHOX2B and GATA3 remained suppressed at day 28 (Fig. 5G), providing insights into the mechanism underlying the durable growth arrest in this treatment group (group V in Fig. 4A). Thus, continued inhibition of KAT6A/B activity leads to sustained low expression levels of MYCN and the adrenergic CRC members PHOX2B and GATA3, which correlates with an increase in Polycomb-mediated silencing, thus explaining the ongoing suppression of tumor-cell proliferation even after cessation of isotretinoin treatment.

### KAT6A/B inhibition plus retinoic acid increases GD2 expression in GD2^low^ neuroblastoma

Anti-GD2 immunotherapy has improved the overall outcome for patients with high-risk neuroblastoma when given as post-consolidation therapy in combination with isotretinoin.^8^ GD2 is a disialoganglioside expressed on the cell surface of most neuroblastoma cells that is targeted in the current treatment of high-risk neuroblastoma using anti-GD2 therapies; however, its expression is often lost at the time of disease recurrence.^39,40–42^ The density of GD2 expression on the surface of neuroblastoma cells likely governs the response to anti-GD2 targeted therapies through antibody-dependent cell-mediated cytotoxicity (ADCC) from NK cells and macrophages; thus, strategies to increase GD2 cell-surface density would be expected to improve treatment response and impart a clinical benefit. To test whether epigenetic reprogramming with isotretinoin plus KAT6A/B inhibition increases the expression of GD2 on the cell surface, the *MYCN*-amplified neuroblastoma cell lines BE2C, NGP and NB-1, which each lack detectable GD2 expression, were treated with either isotretinoin or PF-9363 alone or both drugs in combination. Using flow cytometry, we found that neither treatment alone caused consistent upregulation of GD2 on the cell surface in the tested cell lines. Only the combination of PF-9363 plus isotretinoin caused 10-15-fold increased GD2 expression consistently in all three cell lines (Fig. 6A, B).

**Figure 6:**
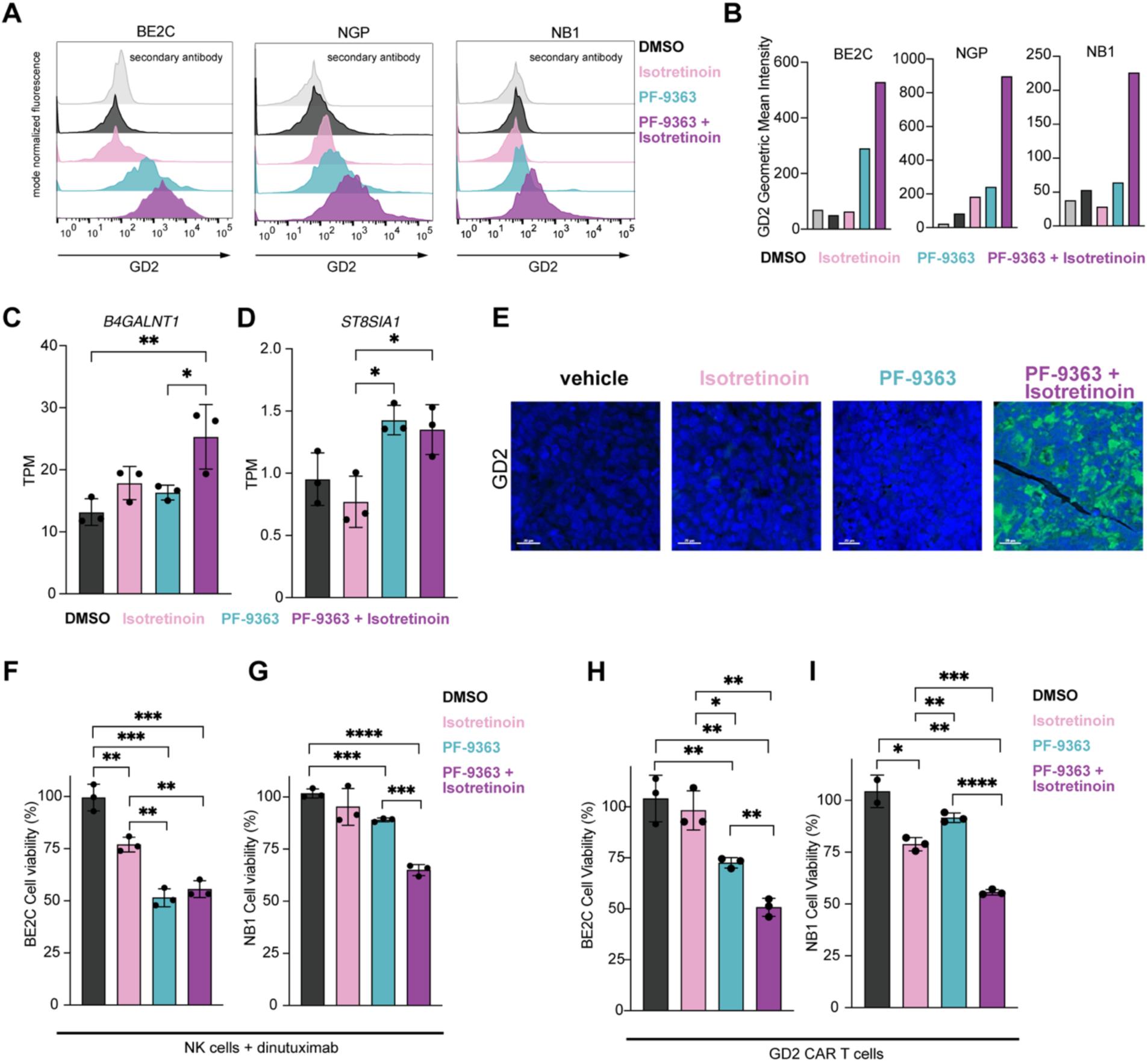
Inhibition of KAT6A/B activity results in enhanced efficacy of GD2 targeting immunotherapy in neuroblastoma. A) Flow cytometry showing GD2 expression levels in the GD2^low^ cell lines BE2C, NGP and NB1 treated for 14 days with DMSO, isotretinoin alone, PF-9363 alone, or isotretinoin plus PF-9363. B) Mean fluorescence intensity for GD2 staining shown in panel A. C-D) Gene expression levels (in TPM from RNA-seq) of *B4GALNT1* (C) and *ST8SIA1* (D) in BE2C cells treated for six days with DMSO, isotretinoin alone, PF-9363 alone, or isotretinoin plus PF-9363. Shown are three biological replicates, and the values represent the mean ± SD. Significance was determined by one-way ANOVA. E) Immunofluorescence for GD2 from FFPE tissue sections of BE2C xenografts in the indicated treatment groups obtained on day 14 of treatment. Scale bar = 20 μm. F-I) Cell viability of BE2C and NB1 cells following both 1) treatment with either DMSO, isotretinoin alone, PF-9363 alone, or isotretinoin plus PF-9363 for 14 days and subsequent 2) co-culture with F-G) NK cells and dinutuximab for 24 hours or H-I) GD2 CAR T cells at an effector-to-target (E:T) ratio of 0.5:1 for 48 hours (*n*=3 samples per treatment group). Data represent mean values ± SD. Significance was determined by one-way ANOVA. *\*p<0.05; **p*<0.001; *\*\*\*p*<0.001; *\*\*\*\*p*<0.0001. Each compound was administered at a concentration of 1 μM.

GD2 disialoganglioside expression on the surface of neuroblastoma cells is derived by a complex synthesis from ceramide mediated by several sialyltransferases and glycosyltransferases.^43^ B4GALNT1 and ST8SIA1 have been implicated as rate-limiting enzymes in this process.^44–46^ Thus, to investigate the mechanisms underlying the upregulation of GD2 expression on the cell surface of neuroblastoma cells, we analyzed the expression levels of *B4GALNT1* (encoding GD2 Synthase) and *ST8SIA1* (encoding GD3 Synthase). Expression of *B4GALNT1* in BE2C cells was unchanged after treatment with either isotretinoin or PF-9363 alone, but was significantly upregulated when treated with the drug combination (Fig. 6C). The expression level of *ST8SIA1* in BE2C cells was generally low and unchanged with isotretinoin treatment, but upregulated when treated with either PF-9363 alone or PF-9363 plus isotretinoin (Fig. 6D). Thus, the combination treatment-mediated an increase in cell surface GD2 expression, which is linked to the transcriptional upregulation of genes required for GD2 synthesis and expression on the cell surface.

To test if GD2 expression is inducible on the surface of neuroblastoma cells not only *in vitro* (Fig. 6A-B) but also *in vivo* through treatment with isotretinoin plus PF-9363, we performed anti-GD2 immunofluorescence on FFPE tissue sections from drug-treated BE2C xenograft tumors. Consistent with our *in vitro* findings, control-treated BE2C tumors did not express GD2 on the cell surface when engrafted in NSG mice (Fig. 6E). Importantly, only combination treatment, but not isotretinoin or PF-9363 alone, induced cell-surface expression of GD2 on tumor cells *in vivo*, similar to the results *in vitro* (Fig. 6E).

To determine whether ADCC is increased following the upregulation of GD2 on the cell surface observed with the combination treatment, we tested whether the ability of NK cells to kill neuroblastoma tumor cells can be enhanced by treatment with either isotretinoin or PF-9363 alone or both in combination when given together with dinutuximab, a monoclonal antibody targeting GD2 used in clinical practice. While treatment with each compound alone decreased the number of viable BE2C cells when co-cultured with NK cells in the presence of dinutuximab, the effect was the largest after drug treatment with PF-9363 or PF-9363 plus isotretinoin (Fig. 6F). By contrast, in NB1 cells, only the double treatment with isotretinoin plus PF-9363, but not the single treatment with isotretinoin led to significant NK-mediated killing of NB1 cells when co-cultured with NK cells and dinutuximab (Fig. 6G).

To assess the effect on specific GD2-mediated tumor cell killing that does not rely on NK cell function and activity, we tested the ability of GD2-specific CAR T cells to kill BE2C and NB1 cells after treatment with either drug alone, or both in combination, compared to DMSO control. Similar to the co-culture with NK cells and dinutuximab, the combination treatment with PF-9363 plus isotretinoin mediated the largest decrease in viable neuroblastoma cells in both cell lines BE2C (Fig. 6H) and NB1 (Fig. 6I) and was more effective than treatment with either compound alone, even at higher ratios of CAR T cells to tumor cells (Supplementary Fig. S5).

In summary, the inhibition of KAT6A/B activity by PF-9363 in combination with isotretinoin has important advantages over isotretinoin alone for the treatment of high-risk neuroblastoma (Fig.7). The combination treatment not only enforces durable growth arrest of tumor cells, but also markedly increases GD2 expression on the neuroblast cell surface and enhances the killing of the neuroblastoma cells by GD2-targeted immunotherapies (Fig. 7). Thus, the incorporation of a KAT6A/B inhibitor into the current combination therapy with isotretinoin plus dinutuximab for high risk neuroblastoma synergizes with both the antiproliferative activity of isotretinoin and the neuroblastoma cell killing activity of dinutuximab to improve the therapy of high risk neuroblastoma.

**Figure 7:**
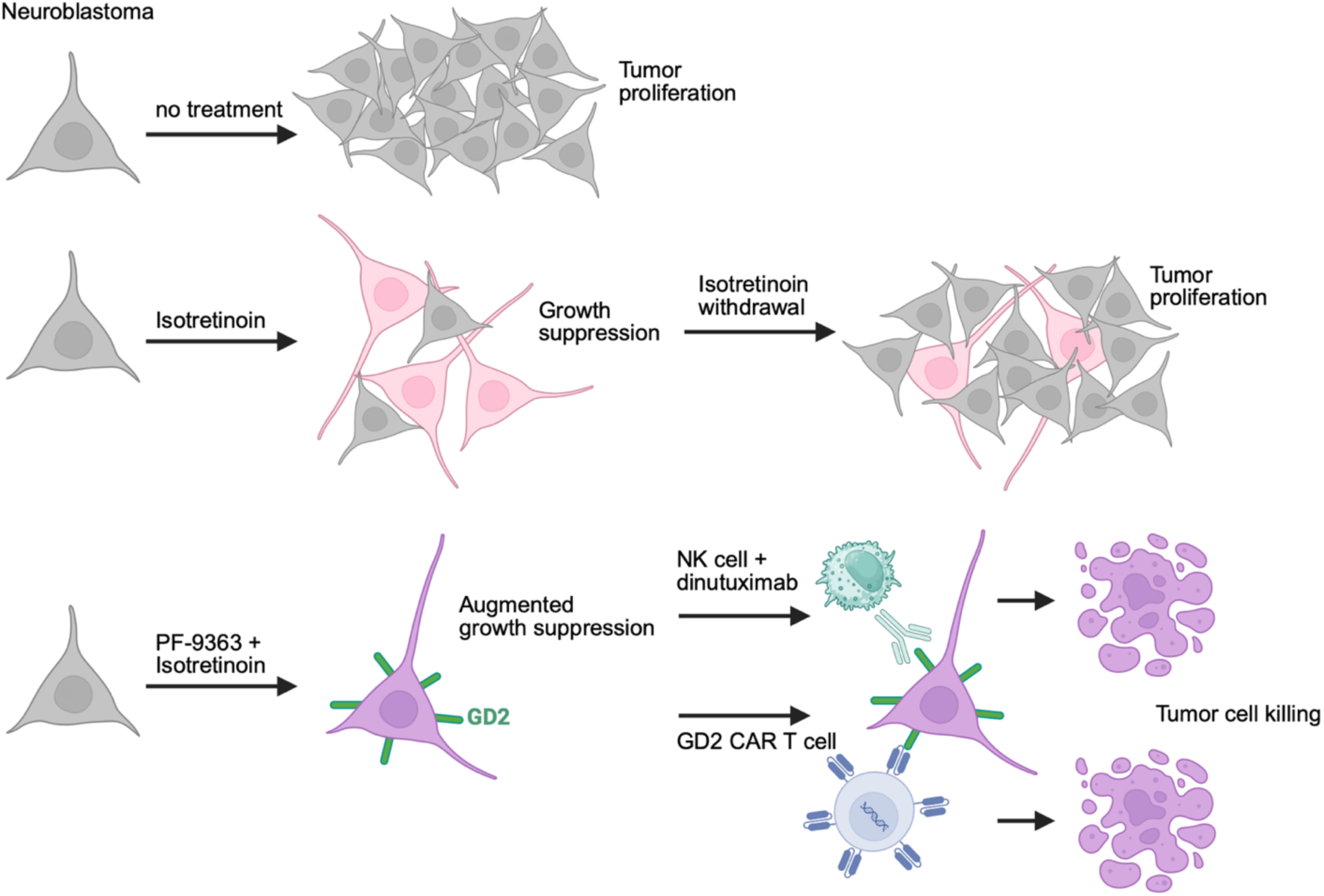
KAT6A/B inhibition enhances the efficacy of retinoic acid and GD2 targeting immunotherapy in neuroblastoma. Schema showing proposed benefit of novel combination treatment using the retinoic acid derivative isotretinoin together with the KAT6 inhibitor PF-9363 to enhance retinoid-induced growth suppression of neuroblastoma, upregulate GD2 expression on the cell surface of GD2^low^ neuroblastoma cells, to increase the efficacy of GD2 targeting immunotherapy - either using the GD2 antibody dinutuximab, or GD2 CAR T cells. The increased efficacy of GD2 targeting immunotherapy has the benefit to induce tumor cell killing rather than only suppressing tumor growth through isotretinoin treatment alone.

## Discussion

Retinoic acid derivatives have potent differentiating and growth-suppressive effects for precancerous states such as leukoplakia or cervical dysplasia,^47–49^ and in a variety of cancers mainly APL and neuroblastoma.^1,2,50–54^ In children with high-risk neuroblastoma, isotretinoin treatment following consolidation therapy improved 3-year event-free survival from 29% to 45%,^2^ but its benefits for long-term overall survival are uncertain.^3^ Here, we sought to improve the antiproliferative effects of retinoic acid in neuroblastoma by identifying a second drug that synergizes with the anti-tumor properties of retinoic acid when both drugs are given in combination, analogous to the very effective combination of ATRA plus arsenic trioxide treatment for APL.^55^ By screening 452 epigenetic inhibitors, we discovered that the KAT6A/B inhibitor PF-9363 synergizes with isotretinoin to inhibit cell growth and augment neuronal differentiation *in vitro* (Figs. 1 and 3). Importantly, our single-cell RNA-seq experiments highlight the limitations of isotretinoin monotherapy in neuroblastoma, demonstrating that isotretinoin is insufficient to drive terminal differentiation for all cells, leaving ∼ 40% of cells in an oligo- or unipotent state, poised to proliferate immediately upon isotretinoin withdrawal (Fig. 3F, G). The combination of isotretinoin and the KAT6A/B inhibitor PF-9363 largely overcomes this limitation and is much more effective in inducing neuronal differentiation (Fig. 3F, G) and synergistically suppresses tumor growth (Fig. 1E).

Moreover, in contrast to single-agent treatment with isotretinoin, in which drug withdrawal leads to rapid tumor proliferation, this novel combination induces durable growth-suppression *in vitro* and *in vivo*, as long as PF-9363 is continued, which is likely due to sustained downregulation of growth-promoting transcription factors including MYCN (Fig. 4 and 5). Importantly, we demonstrate that this new combination treatment induces high levels of GD2 expression on the surface of neuroblastoma cells, which augment GD2-specific tumor-cell killing through ADCC and CAR T cells (Fig. 6F-I).

Currently, post-consolidation treatment for children with high-risk neuroblastoma receiving treatment according to Children’s-Oncology-Group protocols includes six courses of isotretinoin treatment alternating with anti-GD2 antibody therapy. Isotretinoin treatment cycles are limited to 14 days due to toxicity, mainly affecting the skin, which requires a 14-day break before the next cycle can be initiated. The fact that the antiproliferative effects of isotretinoin alone are completely reversible in our *in vitro* and *in vivo* studies (Fig. 4) likely explains the lack of long-term survival benefit of isotretinoin treatment.^3^ Previous attempts to identify novel therapies that enhance the efficacy of isotretinoin focused on its combination with vorinostat, a potent inhibitor of histone deacetylase activity (HDAC).^56,57^ However, the addition of vorinostat to isotretinoin treatment did not demonstrate any benefit for patients with refractory neuroblastoma in a phase-I trial.^58^ In our drug screen, five HDAC inhibitors were among the 28 compounds which led to enhanced growth-suppression of neuroblastoma when combined with retinoic acid (blue bars in Fig.1C). However, the effects of adding HDAC inhibitors to isotretinoin were modest in comparison to the more pronounced effects observed with KAT6A/B inhibitors, including PF-9363 (Fig. 1C-E), which not only synergizes with isotretinoin to inhibit neuroblastoma growth but also enforces sustained growth arrest in combination with isotretinoin (Fig. 4A, E).

Inhibitors of the H3K23 acetylase paralogs KAT6A and KAT6B have shown on-target anti-tumor activity in preclinical studies of several hematological malignancies and solid tumors.^59,60^ The first-in-class KAT6A inhibitor, called WM-1119, has shown to have potent anti-tumor activity in lymphoma and *KAT6A*-rearranged AML *in vitro*.^27,61^ Based on the anti-tumor activity of the second-generation KAT6A/B inhibitor PF-9363 in estrogen-receptor-positive breast cancer in xenograft models,^59,62^ the clinical analog, PF-07248144, is currently under investigation in early phase-I/II trials for patients with heavily pre-treated breast cancer and has demonstrated durable anti-tumor activity in these patients.^28^ In this clinical trial, the daily oral administration of PF-07248144 showed a tolerable safety profile, supporting our reasoning for clinical evaluation of continuous daily oral dosing of a KAT6A/B inhibitor in combination with isotretinoin as post-consolidation treatment for patients with high-risk neuroblastoma. This recommendation is based on our preclinical studies in neuroblastoma xenografts (Fig. 4), which demonstrated the potential of this combination both i) to improve the durability of isotretinoin-induced growth arrest and ii) to upregulate GD2 expression to improve the effectiveness of GD2-targeted therapy (Fig. 6), without evidence of weight loss or other toxicity in murine models.

While anti-GD2 antibody therapy increases the event-free survival of children with high-risk neuroblastoma from 46% to 56%,^8^ its efficacy is limited due to the heterogeneous expression of GD2 in neuroblastoma tumors. Moreover, the apparent ease with which GD2 is downregulated in neuroblastoma cell lines by H3K27me3 accumulation across regulatory regions of GD2 synthetic enzymes,^35^ suggests that this mechanism may be active to generate GD2-resistant variants during treatment of patients. Thus, inducing high levels of GD2 expression on tumor cells through the combination of a KAT6A/B inhibitor plus isotretinoin, as observed in our studies (Fig. 6A, E), may offer additional benefits for patients with GD2^low^ tumors or for those in whom GD2-negative variants emerge and drive relapse during GD2-targeted immunotherapy. Increasing the efficacy of GD2-targeted immunotherapies and enforcing tumor cell-specific killing through either ADCC by NK cells/macrophages via dinutuximab, or CAR T cells (Fig. 6F-I), may be beneficial not only in neuroblastoma but also in other tumor types that express GD2, such as rhabdomyosarcomas, gliomas, melanomas, breast cancers, small-cell lung cancers, Ewing sarcomas and osteosarcomas.

Epigenetic reprogramming with other agents has been studied as a strategy to enforce higher levels of GD2 expression with the goal of increasing the effectiveness of GD2-targeting immunotherapies. Mesenchymal neuroblastoma, a subtype of neuroblastoma arising in disease relapse and associated with chemotherapy resistance, often has low GD2 cell-surface expression levels, and EZH2 inhibitors have been shown to increase GD2 expression and a transition to the adrenergic cell state.^44^ In our studies, even the adrenergic neuroblastoma cell lines BE2C and NGP demonstrate low GD2 expression, which emphasizes that the outgrowth of neuroblastoma cells lacking GD2 expression is of clinical significance in either mesenchymal or adrenergic neuroblastomas and that testing of patients’ tumor samples for GD2 expression at the start of treatment and at the time of disease progression will be important in future clinical trials.

Despite their key roles in regulating neuroblastoma proliferation, therapeutic strategies to directly target MYCN or transcription factors such as PHOX2B and GATA3 have been difficult. Retinoic acid is effective in neuroblastomas because it rewires the enhancer landscape and induces a distinct set of transcription factors that drive a differentiated cell state, leading to decreased expression of MYCN and other adrenergic transcription factors accompanied by growth arrest.^24^ However, upon retinoid withdrawal, PHOX2B, GATA3 and MYCN are rapidly upregulated and the proliferative adrenergic CRC is reactivated, leading to rapid tumor re-growth (Fig.4). By contrast, the addition of a KAT6A/B inhibitor synergizes with isotretinoin to maintain the retinoid-induced repression of adrenergic CRC transcription factors and prevents tumor re-growth when isotretinoin is withdrawn, as long as the KAT6A/B inhibitor is continued (Fig.4 and 5). Acetylation of H3K23 appears to be necessary to reestablish open chromatin needed to re-express high levels of MYCN, PHOX2B and GATA3 after retinoid withdrawal, likely related to its role at sites of active regulatory elements, where it colocalizes with H3K27 acetylation (Fig. 2A-C). Our results suggest that H3K23ac may be required as a precursor to the removal of H3K27me3 repressor histone modifications associated with the regulatory elements of genes encoding adrenergic TFs (Fig. 5). However, the precise mechanisms underlying the role of H3K23 acetylation in forming active regulatory enhancers warrant further investigation.

In summary, our work indicates that altering the epigenetic cell state through the combination of retinoic acid plus KAT6A/B inhibition is likely to be beneficial for treating high-risk neuroblastoma. We show that this strategy alters gene expression in neuroblastoma, leading to sustained growth suppression and induction of GD2 expression on neuroblastoma cells to levels that increase the effectiveness of GD2-targeted immunotherapy. This combined targeted therapy approach therefore merits additional preclinical testing and eventual clinical evaluation in children with high-risk neuroblastoma.

## Materials and Methods

### Cell lines

Cell lines were obtained from the ATCC (BE2C RRID:CVCL_0529) and DSMZ (NGP, LAN-5, KELLY). NB1 was obtained from the Stegmaier laboratory at the Dana-Farber Cancer Institute, Boston. All cell lines were short-tandem-repeat (STR)–tested for identity prior to use. Human neuroblastoma cell lines were cultured at 5% CO2 in RPMI medium containing 10% fetal bovine serum and 1% penicillin-streptomycin and were routinely validated to be free of Mycoplasma species.

### Drug screen

5000 BE2C cells were plated in each well of a white 96-well plate in 100 μl of total medium containing either DMSO or 5 μM ATRA alone or in combination with one of the 452 epigenetic-modifying compounds at a concentration of 1 μM (see Supplementary Table S1). Cells were incubated for five days, and then cell viability was assayed with CellTiter-Glo (Promega) according to the manufacturer’s protocol. Calculation of z-score: The z-score is defined as the relative growth of cells treated with compound alone minus the average growth of all treated cells in the screen divided by the standard deviation. The relative growth equals the ratio of viable-cell number treated with compound or compound plus isotretinoin versus the viable-cell number treated with DMSO control.

### Proliferation assays

For cell-proliferation assays, neuroblastoma cells were plated on 24-well (10,000 cells/well) or 6-well (20,000 cells/well) dishes in RPMI medium containing DMSO, 1 μM isotretinoin, 1 μM PF-9363 or 1 μM isotretinoin plus 1 μM PF-9363. Cell viability was assessed at day seven (unless noted otherwise) by counting live cells using the Countess 3 FL Automated Cell Counter (Invitrogen). For long-term proliferation assays including cell counts on days 7, 14, 21 or 28, neuroblastoma cells were plated on 100-mm dishes (100,000 cells/well) and maintained in RPMI medium containing either DMSO (I), 1 μM isotretinoin (II), 1 μM PF-9363 (III) or 1 μM isotretinoin plus 1 μM PF-9363 (IV). For treatment groups I and III, cells were plated into a single dish and harvested on respective days, and viable-cell count was documented. 1x10e6 cells from each treatment group were re-plated and kept in RPMI medium containing either DMSO (I) or PF-9363 (III) until the next collection time point. For the treatment groups II and IV, cells were plated into four dishes at the beginning of the experiment, and at each data point, one dish of each group was collected, and the viable-cell count documented.

### Chemicals

The drug library (HY-LD-000001689), isotretinoin and PF-9363 were obtained from MedChemExpress. ATRA was obtained from Sigma-Aldrich. Cell-culture–grade DMSO was purchased from ATCC. Compounds were resuspended in DMSO to a stock concentration of 10 mM and added directly to cell-culture medium at the indicated concentrations.

### Animals

Eight-week-old female NOD.Cg-*Prkdc*scid *Il2rg*tm1Wjl/SzJ RRID:IMSR_JAX:025216 (NSG) mice (The Jackson Laboratory) were used for tumor xenograft studies and MTD testing.

### Plasmids and lentiviral transduction

Stable *Cas9:sgRNA*-expressing cell lines were created using lentivirus produced in 293T cells (RRID:CVCL_0063). Briefly, sgRNA target sequences (Table S2) were cloned in the lentiCRISPRv2 vector (Addgene plasmid #52961), as previously reported.^24,63^ Plasmids were transfected using X-tremeGENE (Roche) along with pMD2.G (RRID:Addgene_12259) and psPAX2 (RRID:Addgene_12260) to generate viral particles. mCherry-luciferase driven by EF1α promoter lentiviral particles were generated as previously described.^64^ BE2C and NB1 cells were transduced with virus in the presence of 10 µg/ml protamine sulfate and were selected with puromycin (1 µg/ml) and expanded before evaluation.

### Immunohistochemistry

Formalin-fixed, paraffin-embedded (FFPE) blocks were prepared and sectioned, and immunohistochemistry was conducted, at the Research Pathology Core lab at the Dana-Farber Cancer Institute. Ki67 expression was detected using a primary antibody (detailed in Table S3) and visualized with a diaminobenzidine-peroxidase system (EnVision+, Dako). Counterstaining was performed with Mayer’s hematoxylin, and slides were imaged using the Echo Revolve4 inverted-microscopy system. GD2 immunofluorescence was detected following a previously described protocol.^65^ Imaging of the tissue samples was performed using the Akoya Biosciences PhenoCycler®-Fusion 1.0 system, with subsequent visualization using Phenochart and inForm (Akoya AI-based software).

### Western blotting

Whole-cell lysates were obtained from cells growing in culture using radioimmunoprecipitation-assay buffer containing protease and phosphatase inhibitors (Cell Signaling Technology). Histone extracts (HE) were prepared with the Total Histone Extraction kit from Abcam (ab113476) according to the manufacturer’s protocol. Briefly, lysates/HE were quantified by Bradford assay (Bio-Rad), and 10 μg of extracted protein (lysates) or 3 μg of HE was separated using Novex SDS–polyacrylamide-gel-electrophoresis reagents and transferred to nitrocellulose membranes (Life Technologies). Membranes were blocked in 5% milk protein and incubated with primary antibodies (Table S3) overnight followed by secondary horseradish-peroxidase–linked goat anti-rabbit and anti-mouse (Cell Signaling Technology) antibodies (1:1000) according to the manufacturers’ instructions. Antibody-bound membranes were incubated with SuperSignal West Pico chemiluminescent substrate (Thermo Fisher Scientific) and developed using HyBlot CL autoradiography film (Thomas Scientific).

### Flow cytometry detection of GD2

Cell lines were trypsinized into suspension and washed twice with PBS + 2% FCS before staining with primary antibody (see Table S2), or only secondary antibody as a control. Antibodies were incubated for 60 min at room temperature. Cells were filtered through a 40-µM filter and immediately measured on a BD Celestra or Fortessa flow cytometer. Further flow cytometry analysis and data analysis was performed as previously reported.^44^

### Spike-in normalized RNA-seq

BE2C sg*NT1*, sg*KAT6A* and sg*KAT6B* were grown in triplicate using six-well plates and collected directly into TRIzol. DMSO-, isotretinoin-, PF-9363- and isotretinoin plus PF-9363-treated BE2C cells were grown in triplicate using six-well plates and collected directly into TRIzol. Each compound was administered at 1 µM. ERCC spike-in RNA (Life Technologies) was diluted 1:10 in nuclease-free water and added directly to TRIzol lysates after being normalized to cell number as previously described.^66^ Library preparation and sequencing were conducted as previously described.^66^ Graphs were created using GraphPad Prism, RRID:SCR_002798 10.2.2.

RNA expression was quantified largely as in Weichert-Leahey et al.^66^ A reference genome containing the sequences of the hg19 version of the human reference genome and the sequences of the ERCC spike-in probes was constructed. Raw sequencing reads were aligned to this reference genome using hisat v2.1.0^67^ in paired-end mode with default parameters using all input FASTQs for each sample. Output SAM files were converted to BAM and sorted by name using samtools^68^ v1.9. A reference gene annotation was created by combining the positions of RefSeq genes downloaded 7/5/2017 and positions of the artificial ERCC chromosomes. Expression of the genes in this annotation was calculated using htseq^69^-count v0.11.2 with parameters -m intersection-strict -I gene_id –stranded-reverse. Per-gene transcripts-per-million (TPM)^69^ values were computed by first calculating the number of unique base-pairs in all exons of all isoforms connected to the same gene ID, then calculating a per-sample normalization term = sum of (readcount * readlength/exon base-pairs) for all genes, then calculating TPM for each gene as readcount * readlength/exon base-pairs * 1e6/normalization term.

Cell number normalization of TPMs was performed as previously,^66^ using all samples within a batch of experiments, i.e., all BE2C drug-treated samples comprised a batch, and all BE2C knockdown samples comprised a separate batch. Briefly, TPM values between 0 and 0.1 were set to 0.1, and a pseudocount of 0.1 was added to all values. The TPM values of the ERCC probes were used to normalize other calculated TPMs using loess.normalize from the affy R package. Normalized TPMs were averaged across all replicates of the experiment. Log2 fold-changes of these averages between condition-pairs were calculated after adding a pseudocount of 1.

For differential expression analysis, cell-number-normalized TPM values were used. Genes were considered expressed if they had an average TPM>1. A log2 TPM post-normalization fold-change of 0.5 was considered significant in genome-wide. A one-way ANOVA test was used to compare TPM levels of individual genes using triplicates for each cell-number-normalized condition. Significantly upregulated genes in both sg*KAT6A* and sg*KAT6B* versus sg*NT1* BE2C cells were input into the metascape database (RRID:SCR_016620)^70^ and a customized enrichment analysis was performed including the following GO terms: GO Molecular Functions, GO Biological Processes and GO Cellular Components.

### Single-cell RNA-seq analysis

#### Initial pre-processing and quality control filtering of scRNA-seq data

Read count matrices for scRNA-Seq data were generated from raw FASTQ files using Cell Ranger (version 7.1.0). Reads were aligned to the GRCh38 transcriptome reference (version refdata-gex-GRCh38-2020-A). The resulting gene count matrices were processed and analyzed in the R programming language (version 4.4.0) using the Seurat package.^71^ Ambient RNA was removed using the SoupX package^72^ (version 1.6.2) using the *autoEstCont* and *adjustCounts* functions with default parameters. Quality control filtering was applied to each cell, using filters of 3000 < nFeature_RNA < 8000 and mitochondrial read percentage < 20%. Cell doublets were called and removed using the scDblFinger package^73^ (version 1.18.0) with default parameters.

### Normalization, clustering, cell cycle scoring, and potency analysis of scRNA-seq data

Normalization and analysis of scRNA-Seq data was performed using the Seurat package (version 5.1.0). Cell cycle scores were computed using the *CellCycleScoring* method of Seurat with annotated cell cycle genes (2019 update). Each experimental treatment sample was normalized by *SCTransform* (version 2) with regression of the cell cycle difference score (S Score minus G2M Score). This workflow was previously found to regress out less interesting differences in cell cycle phase among proliferating cells while better preserving cellular differentiation signals for downstream clustering. The resulting experiments were then merged without integration. Principal component analysis (PCA) was performed on the merged Seurat object and *FindNeighbors* and *RunUMAP* were run using the top 30 PCs for Uniform Manifold Approximation and Projection (UMAP) dimensionality reduction. Louvain clustering was performed on the merged experiment object using *FindClusters* at a resolution of 0.3 and using the top 30 PCs. Cellular potency was called on the merged raw count data using the CytoTRACE2 package^38^ (version 1.0.0), specifying *is_seurat=TRUE* and *species=”human”* and otherwise using default parameters. Visualization was performed using the SCpubr package^74^ (version 2.0.2).

### CUT&RUN

CUT&RUN coupled with high-throughput DNA sequencing was performed using antibodies listed in Table S3 and Cutana pA/G-MNase (EpiCypher) according to the manufacturer’s protocol as previously described.^24^

### CUT&RUN, ATAC-Seq, and ChIP-Seq processing

Irrespective of whether the sample itself contained spike-ins, raw ChIP-Seq or CUT&RUN reads were first aligned to the E. coli reference genome (Escherichia_coli_K_12_MG1655) using bowtie (RRID:SCR_005476) v1.2.2 with parameter -k 1 in single-end mode and -l set to read length. Unmapped reads were then aligned to the human reference genome revision hg19 using bowtie with parameters -k 1 -m 1 –best in single-end mode with -l set to read length, and the outputs were converted to BAM using samtools v1.9. Coverage tracks showing read pileup were created using MACS v1.4.1 using parameters -S -w –space=50 –nomodel –shiftsize=200, normalized by the millions of mapped reads, converted to bigwig format and visualized in the Integrative Genomics Viewer (IGV) browser version 2.16.1.^75^

Peaks of H3K27ac were identified using MACS v1.4.1 with corresponding input control and -p 1e-9. Peaks of KAT6A and KAT6B were defined using MACS v1.4.1 with default parameters. To determine the distribution of KAT6A and KAT6B sites genome-wide, promoters were defined as 4kb regions centered on the transcription start sites of each transcript from the gene list described above, and enhancers were defined as H3K27ac peaks that did not overlap these promoters. H3K27ac peaks were defined using MACS v1.4.1 with -p 1e-9. Overlaps were determined using bedtools intersect [CITE PMID: 20110278]. 4kb windows centered on transcription start sites of RefSeq genes were generated and used as input for bamToGFF (https://github.com/BradnerLab/pipeline) with parameters -m 50 -r to generate matrices with reads-per-million normalized and duplicate-removed coverage of 50 bins apiece for each window. Coverage heatmaps were generated using heatmap.3 ranking each row according to the row sum of the specified factor. Metagenes were created using the same matrices but instead representing the mean of each genomic bin/column.

For quantitative coverage analysis of H3K27me3, presumed PCR duplicate reads were removed using samtools markdup -r. Islands of significant coverage of H3K27me3 reads were separately determined for each sample using SICER (https://github.com/zanglab/SICER2)^76^ with parameters -rt 1 -w 200 -f 76 -g 400 -e 100. These islands were collapsed using bedtools merge for a unified set of regions-of-interest. Islands were assigned to the single gene whose transcription start site was nearest the center of the island using bedtools closest -t first. The duplicate-removed read coverage of each merged island for each sample was calculated using bedtools intersect -c. Read counts for each sample were normalized by the millions of mapped, duplicate-removed reads.

Raw ATAC-Seq reads were analyzed as previously.^11^ Briefly, reads were aligned to the hg19 revision of the human reference genome using bowtie v1.2.2 with parameters -l 75 -k 1 -m 1 – best. Peaks were identified using MACS v1.4.1 with -p 1e-9.

### ChIP-seq

ChIP-seq was performed as previously described.^11^ The antibodies used for each experiment are listed in Table S3. For each ChIP, 5 μg of antibody coupled to 2 μg of magnetic Dynabeads (Life Technologies) was added to 3 ml of sonicated nuclear extract from formaldehyde-fixed cells. Chromatin was immunoprecipitated overnight, cross-links were reversed, and DNA was purified by precipitation with phenol:chloroform:isoamyl alcohol. DNA pellets were resuspended in 25 μl of TE buffer. Illumina sequencing, library construction, and ChIP-seq analysis methods were previously described.

### In Vivo *Studies*

All mouse experiments were performed according to protocols approved by the Dana–Farber Cancer Institute Animal Care and Use Committee (IACUC), and animals were maintained according to institutional guidelines. For xenograft experiments, 8-week-old female NSG mice (The Jackson Laboratory) were subcutaneously implanted with 2.5 × 10^6^ BE2C cells in 50% Matrigel/PBS. Mice were randomly assigned to the following six groups: i) vehicle (*n* = 9) for 28 days, ii) isotretinoin at 50 mg/kg/day on days 1-14 followed by vehicle on days 15-28 (*n* = 9), iii) PF-9363 at 5 mg/kg/day for 28 days (*n* = 5), iv) PF-9363 at 5 mg/kg/day on days 1-14 followed by vehicle on days 15-28 (*n* = 4), v) PF-9363 at 5 mg/kg/day on days 1-28 plus isotretinoin at 50 mg/kg/day on days 1-14 (*n* = 5), and vi) PF-9363 at 5 mg/kg/day on days 1-14 plus isotretinoin at 50 mg/kg/day on days 1-14 followed by DMSO on days 15-28 (*n* = 4). All compounds were given daily by oral gavage. Treatments started four days after BE2C-cell injection. Tumors were measured by calipers, and mice were weighed every three days. Animals were euthanized according to institutional guidelines when tumors reached 2,000 mm in length or width, or if animals became moribund. Tumor sizes were compared by Mann-Whitney test. For IHC analysis, NSG mice were xenografted with BE2C and received treatment in groups i-vi and three animals from each treatment group were sacrificed at days 14 or 28.

### CAR T cell production and CAR T cell cytotoxicity assay

Blood from healthy donors (BCH IRB P00032515) was collected into heparin containing tubes. PBMCs were isolated by Ficoll gradient followed by T cell selection using the Pan T Cell Isolation Kit (Miltenyi Biotec). T cells were activated and expanded using CD3/CD28 activation beads in X-vivo media containing 10% FBS, penicillin/streptomycin, glutamax and 50 U/mL IL-2. After 48 hours, T cells were transduced with VSV-typed EF1α-driven GD2 CAR or CD19 CAR lentivirus and expanded for an additional 7-10 days. Tumor cell lines were pre-treated as indicated for 14 days. Tumor cells were replated at 10,000 cells/well, followed by addition of CAR T cells at effector-to-target ratio of 0.25:1, 0.5:1, and 1:1. After 48h, Bright-Glo Luciferase (Promega) was added at 1:1 by volume, and incubated for 2 mins prior to quantification using an EnVision microplate reader (Perkin Elmer). GD2 CAR T cell specific killing was calculated as a fraction from maximum luciferase signal in tumor cells co-cultured with CD19 CAR T cells to account for non-specific background killing.

### Dinutuximab assay

PBMCs were isolated as described above followed by NK cell selection using the EasySep Human NK Cell Isolation Kit (Stemcell technologies) and cultured in NK NK MACS Media (Miltenyi Biotec) supplemented with 500 U/mL recombinant human IL-2 (R&D systems) and 1% penicillin/streptomycin/glutamine. NK cells were cultured for 1-2 days prior to assays. BE2C and NB1 cells were treated with DMSO (I), 1 μM isotretinoin (II), 1 μM PF-9363 (III) or 1 μM isotretinoin plus 1 μM PF-9363 (IV) for 14 days. On day 14 of treatment, cells were collected and plated/ into 96-well plate with 20,000 cells per well. Wells received one of four treatment cohorts in triplicate (24 h): (1) human isotype antibody (1 μg ml−1), (2) dinutuximab (1 μg ml−1), (3) human isotype (1 μg ml−1) with 1:2 (NK/tumor cell) ratio or (4) dinutuximab (1 μg ml−1) with 1:2 (NK/tumor cell) ratio. After the incubation period, Bright-Glo Luciferase (Promega) was added at 1:1 by volume, and incubated for 2 mins prior to quantification using an EnVision microplate reader (Perkin Elmer). Background killing from NK cells alone was subtracted from detected values, and treatments were normalized to cell viability in group 1.

#### Histone extraction and LC-MS/MS analysis

The histones were extracted from BE2C cells treated with either DMSO (I), 1 μM isotretinoin (II), 1 μM PF-9363 (III) or 1 μM isotretinoin plus 1 μM PF-9363 (IV) for 14 days and prepared for chemical derivatization and digestion as described previously.^77,78^ In brief, the lysine residues from histones were derivatized with the propionylation reagent (1:2 reagent:sample ratio) containing acetonitrile and propionic anhydride (3:1), and the solution pH was adjusted to 8.0 using ammonium hydroxide. The propionylation was performed twice, and the samples were dried on speed vac. The derivatized histones were then digested with trypsin at a 1:50 ratio (wt/wt) in 50 mM ammonium bicarbonate buffer at 37°C overnight. The N-termini of histone peptides were derivatized with the propionylation reagent twice and dried on speed vac. The peptides were desalted with the self-packed C18 stage tip. The purified peptides were then dried and reconstituted in 0.1% formic acid. An LC-MS/MS system consisting of a Vanquish Neo UHPLC coupled to an Orbitrap Exploris 240 or Ascend (Thermo Scientific) was used for peptideanalysis. Histone peptide samples were maintained at 7 °C on a sample tray in LC. Separation of peptides was carried out onan Easy-Spray™ PepMap™ Neo nano-column (2 µm, C18, 75 µm X 150 mm) at room temperature with a mobile phase. The chromatography conditions consisted of a linear gradient from 2 to 32% solvent B (0.1% formic acid in 100% acetonitrile) insolvent A (0.1% formic acid in water) over 48 min and then 42 to 98% solvent B over 12 min at a flow rate of 300 nL/min. The mass spectrometer was programmed for data-independent acquisition (DIA). One acquisition cycle consisted of a full MS scan and 35 DIA MS/MS scans of 24 m/z isolation width starting from 295 m/z to 1100 m/z. Typically, full MS scans were acquired in the Orbitrap mass analyzer across 290–1200 m/z at a resolution of 70,000 or 120,000 in positive profile mode with an injection time of 50 ms and an AGC target of 1e6 or 200%. MS/MS data from HCD fragmentation was collected in the Orbitrap. These scans typically used an NCE of 30 or 25, an AGC target of 1000%, and a maximum injection time of 60 ms. Histone MS data were analyzed with EpiProfile.^79^

### Statistical analysis

Data from ChIP-seq, CUT&RUN-seq, and RNA-seq were analyzed as described previously. For statistical comparisons among multiple groups, we used one-way ANOVA with *post-hoc* Tukey test unless otherwise noted. Tumor volumes from animal experiments were analyzed by Mann-Whitney test. Data were analyzed with GraphPad Prism, RRID:SCR_002798 10.2.2., and all error bars represent standard deviation (s.d.) unless otherwise noted. Statistical significance was defined as *P* < 0.05 for all experiments.

## Acknowledgments

We thank Dr. Richard A. Young (Whitehead Institute, MIT), and Drs. Scott Armstrong and Steven DuBois (DFCI) for critical discussions. We thank Dr. Prafulla Gokhale and the Animal Resources Facility/Lurie Family Imaging Center at Dana-Farber Cancer Institute for assistance with animal studies. We thank the Research Pathology Core lab at the Dana-Farber Cancer Institute and Dr. Namrata Singh for assistance with IHC studies.

This work was supported by grants from the National Cancer Institute, National Institute of Health, R35 CA210064 (ATL), R35 CA283977 (KS), K08 CA245251 (ADD), K12 HD052896 (FAC), and T32 HL007574-39 (NWL). This work was also supported by the St. Jude Children’s Research Hospital Collaborative Research Consortium on Chromatin Regulation in Pediatric Cancer (ATL, KS, and BJA) and the American Lebanese Syrian Associated Charities (BJA, ADD) and by a grant from Cookies for Kids’ Cancer (ATL). Additional funding was provided by the Parker Institute for Cancer Immunotherapy and V Foundation for Cancer Research, co-sponsoring a Parker Bridge Fellow Award (DAO). NWL is supported by the Damon Runyon Cancer Research Foundation, the Rally Foundation for Childhood Cancer Research and Hyundai Hopes on Wheels Foundation.

Raw and processed data files will be deposited to the NCBI GEO server under a super-series and summarized in Table S4.

We thank Cicely Jette for helpful editorial comments on the manuscript.

## Authors’ Contributions

**N. Weichert-Leahey:** Conceptualization, resources, data curation, formal analysis, supervision, funding acquisition, validation, investigation, visualization, methodology, writing–original draft, writing–review and editing, project administration. **A. Berezovskaya:** Resources, validation, investigation, methodology. **M.W. Zimmerman:** Conceptualization, formal analysis, investigation, writing–review and editing. **Francesca Alvarez-Calderon:** Formal analysis, investigation, validation, methodology, writing–review and editing. **Marlana Winschel**: Resources, investigation. **Silvi Salhotra**: Resources, Formal analysis, investigation. **Nathaniel Mabe:** Formal analysis, investigation. **Antonio Perez-Atayde:** Formal analysis, investigation, validation. **Francisca N. de L. Vitorino**: Formal analysis, investigation, validation. **Benjamin Garcia:** Supervision. **Ulrike Gerdemann:** Formal analysis, supervision, investigation, writing–review and editing. **Adam D. Durbin:** Conceptualization, supervision, writing–review and editing. **K. Stegmaier:** Conceptualization, resources, supervision, funding acquisition, writing–review and editing. **D. A. Oldridge:** Formal analysis, visualization, methodology, writing-review and editing. **B.J. Abraham:** Conceptualization, resources, data curation, software, formal analysis, validation, investigation, methodology, writing–review and editing. **A.T. Look:** Conceptualization, resources, supervision, funding acquisition, project administration, writing–review and editing.

## Statement of significance

Retinoic acid inhibits growth of neuroblastoma, but its effects are completely reversible upon treatment cessation. Here, we identify PF-9363, an inhibitor of the histone acetyltransferases KAT6A/B, which synergizes with retinoic acid to induce durable growth arrest that persists beyond retinoid withdrawal, while increasing the effectiveness of GD2-targeted immunotherapy.

