## Supplemental Figures and Tables for "KAT6A/B inhibition synergizes with retinoic acid and enhances the efficacy of GD2-targeted immunotherapy in neuroblastoma"

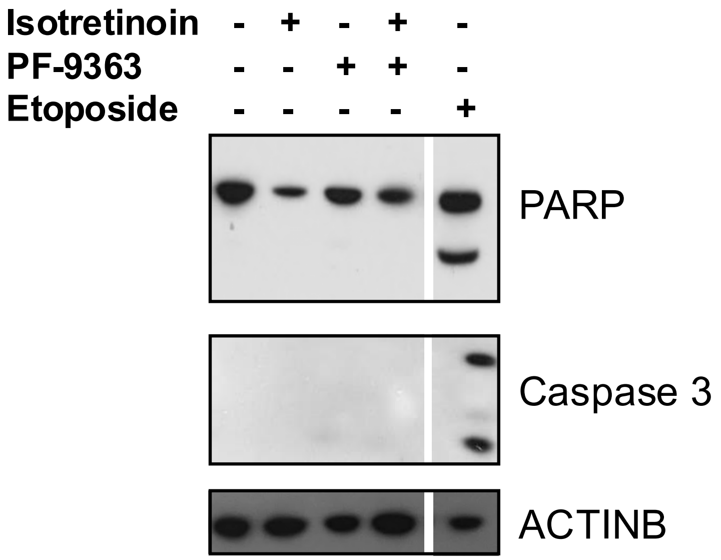


**Supplementary Figure 1: Isotretinoin plus KAT6A/B inhibition does not induce apoptosis in BE2C cells.** Western blotting with cell lysates extracted from BE2C cells treated with the indicated compounds at a concentration of 1 μM each for eight days. Etoposide treatment was administered to BE2C cells for 48 hours as a positive control for the induction of apoptosis. Antibodies against PARP and Caspase 3 were used to detect cleavage events that occur during apoptosis. Beta-actin (ACTINB) was included as a loading control.

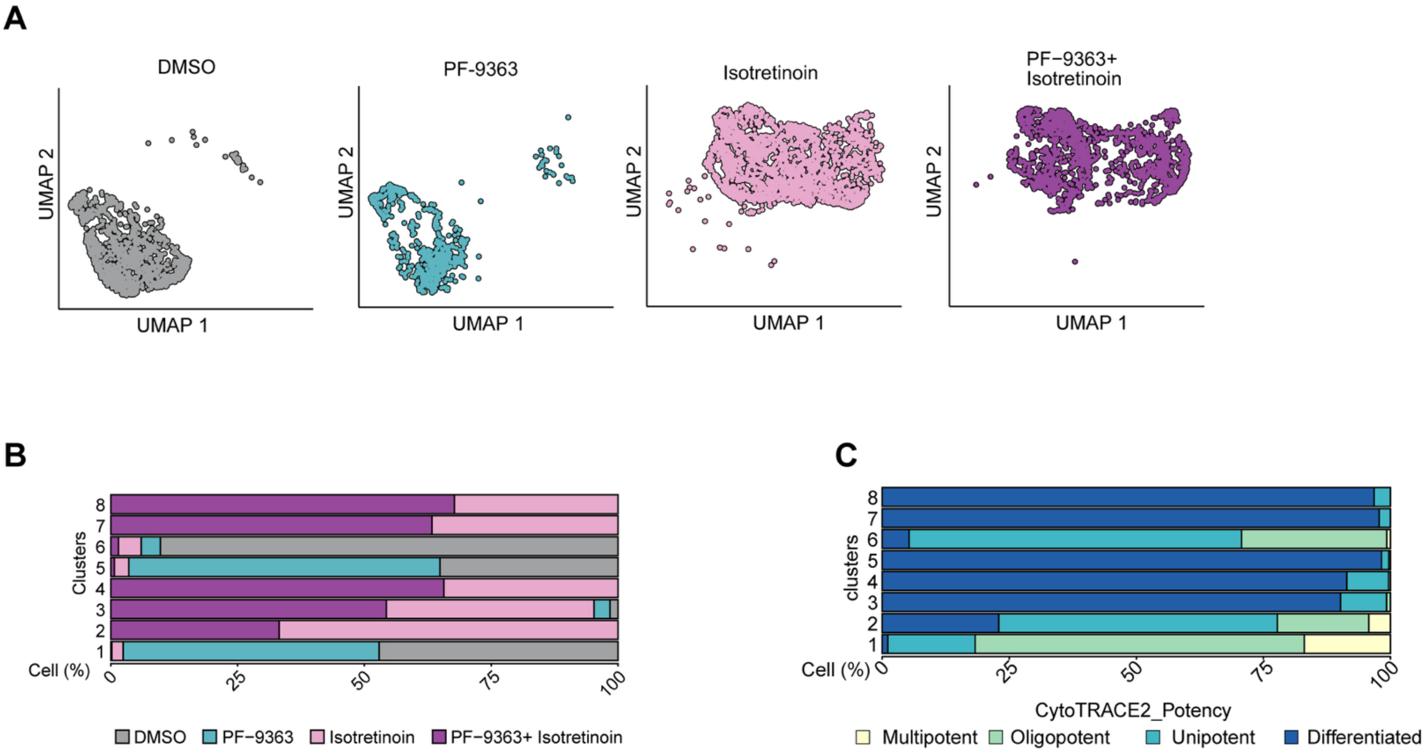


**Supplementary Figure 2: Single-cell RNA-seq analysis identifies augmentation of retinoid-induced differentiation of neuroblastoma through inhibition of KAT6A/B activity.** Single-cell RNA-seq was performed from BE2C cells treated with either DMSO, PF-9363 alone, isotretinoin alone, or isotretinoin plus PF-9363. Each compound was administered at a concentration of 1 μM. UMAP analysis was conducted across all treatment groups merged. A) UMAP analysis from BE2C cells highlighting each treatment condition separately. B) Cluster analysis was performed across all treatment groups merged. Bar chart of each cluster with % cells from each of the four treatment conditions. C) Bar chart of each cluster with % cells assigned to the four categories of the CytoTRACE 2 potency score. Each treatment group is normalized by cell count.

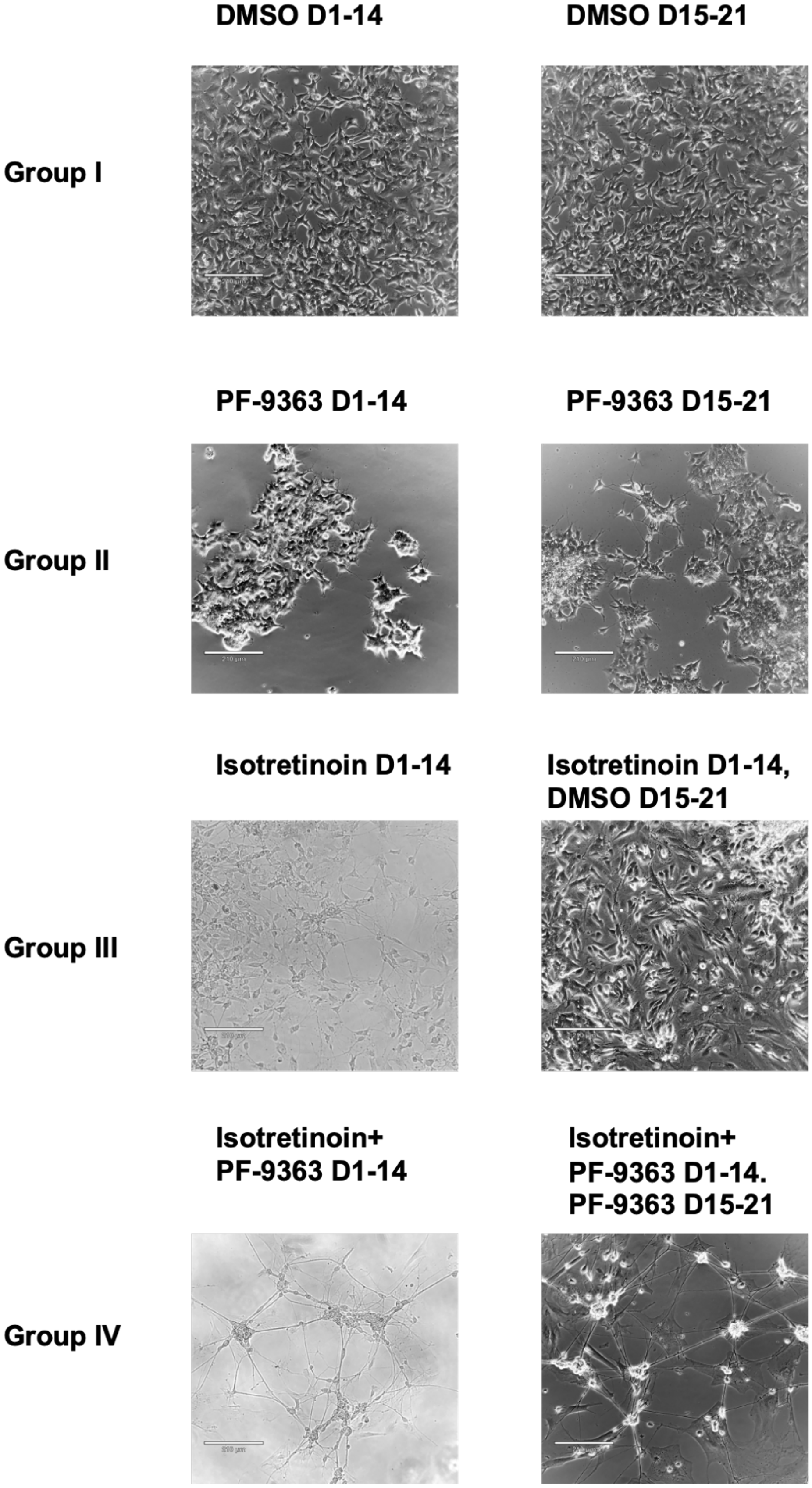


**Supplementary Figure 3: The combination of KAT6A/B inhibition and isotretinoin induces sustained differentiation of neuroblastoma cells.** BE2C neuroblastoma cells were treated for 21 days with DMSO (group I), 21 days with PF-9363 (group II), 14 days with isotretinoin followed by 7 days with DMSO (group III), or isotretinoin plus PF-9363 for 14 days followed by 7 days with PF-9363 (group IV). Morphology of treated BE2C cells was assessed by darkfield microscopy after 14 days of treatment (left column) and at day 21 (right column). Each compound was administered at a concentration of 1 µM. Scale bar = 210 µM.

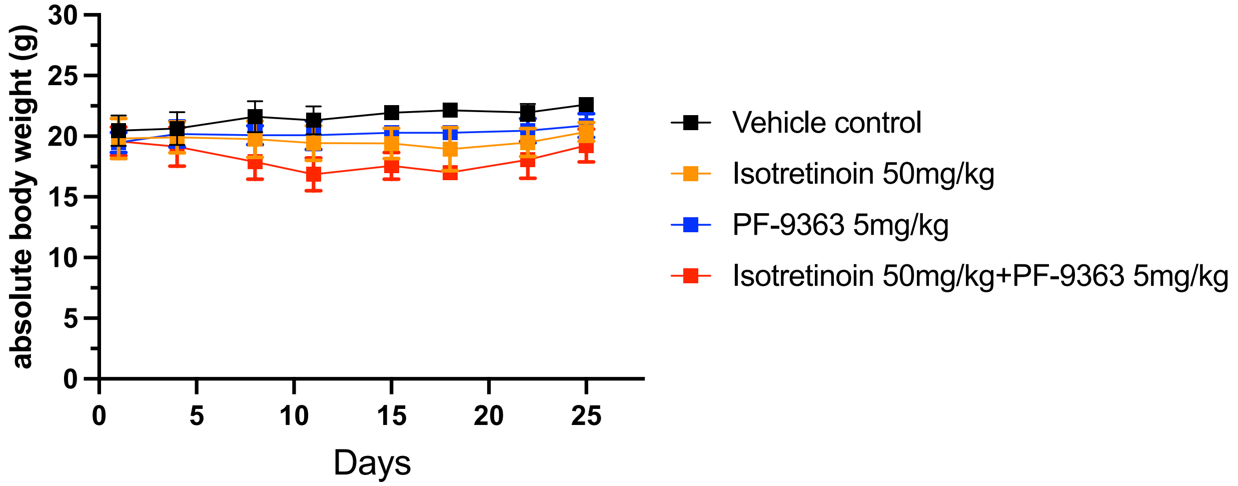


**Supplementary Figure 4: KAT6A/B inhibition in combination with isotretinoin is non-toxic *in vivo.*** Body weight of BE2C xenografts in NSG mice that received treatment with i) either vehicle control for 28 days (black line), ii) isotretinoin at 50 mg/kg/d po daily for 14 days, followed by vehicle control of 14 days (orange line), or iii) PF-9363 5mg/kg/d po alone for 28 days (blue line) or iv) the combination of isotretinoin 50 mg/kg/d po plus PF-9363 5mg/kg/d po for 14 days, followed by either PF-9363 alone or vehicle for 14 days (red line). None of the mice in this experiment lost > 20% of their body weight, which would have required interruption of treatment.

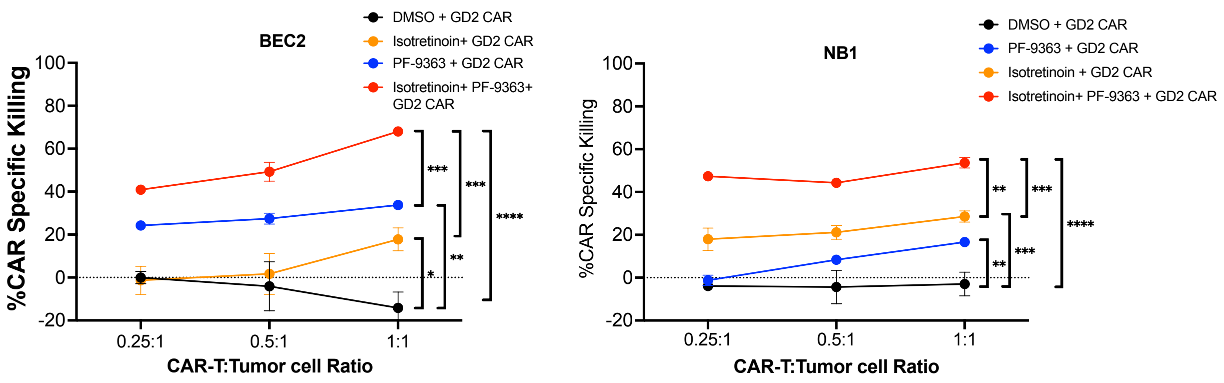


**Supplementary Figure 5: Inhibition of KAT6 activity results in increased effectiveness of GD2 CAR T cell-mediated killing of neuroblastoma cells.** BE2C and NB1 cell killing after treatment with either DMSO (black), isotretinoin alone (orange), PF-9363 alone (blue), or the combination of isotretinoin plus PF-9363 (red) for 14 days followed by co-culture with GD2 CAR T cells at an E:T ratio ranging from 0.25:1 to 1:1 for 48 hours (*n*=3 samples per treatment group). Data are shown as mean+/- SD. Significance was determined by one-way ANOVA. * *p<0.05, **p<0.001, ** p<0.0001*. All compounds administered at a concentration of 1μM each.

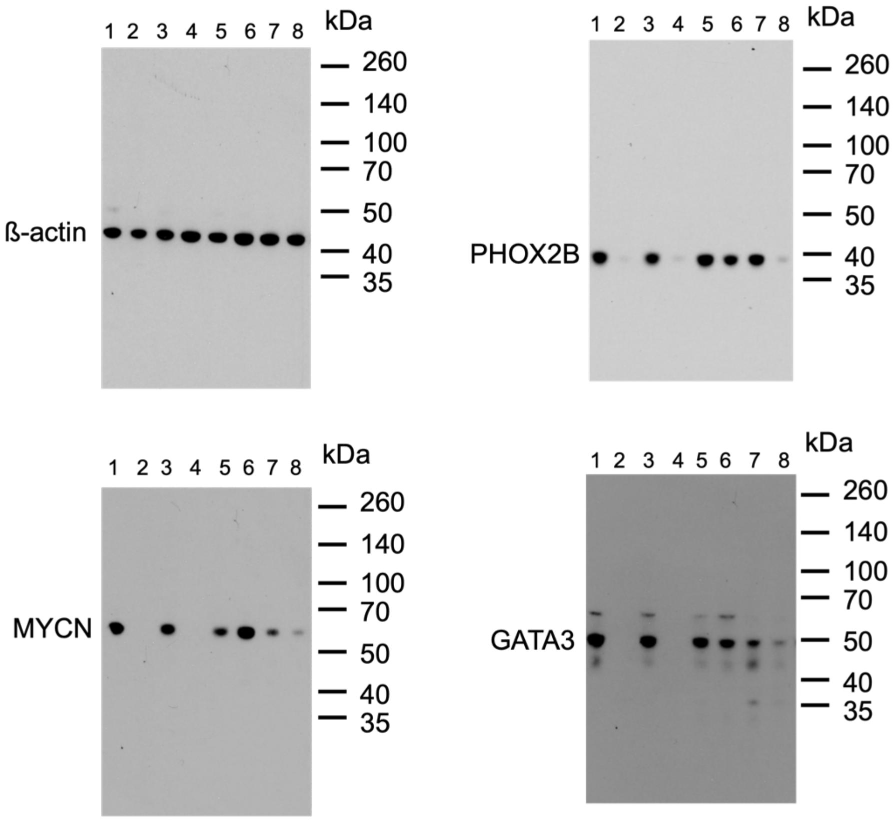


**Supplementary Figure 6:** Representative uncropped western blots for Figure 3F-I: Western blot assay using antibodies against beta actin, PHOX2B, MYCN and GATA3 using cell lysates from BE2C cells treated with 1) DMSO, 2) isotretinoin, 3) PF-9363, 4) PF-9363 plus isotretinoin for 14 days and 5) DMSO for 28 days, 6) isotretinoin for 14 days, followed by 14 days of DMSO, 7) PF-9363 for 28 days and 8) PF-9363 plus isotretinoin for 14 days, followed by PF-9363 only for 14 days.

| |
|---|
| Panobinostat |
| Inauhzin |
| Niraparib tosylate |
| Nicotinamide |
| Chelerythrine Chloride |
| PFI-4 |
| Tranylcypromine (hemisulfate) |
| Phenformin (hydrochloride) |
| [6]-Gingerol |
| MS023 |
| STO-609 |
| GSK-J2 |
| Daminozide |
| BI-847325 |
| NMS-P118 |
| LLY-507 |
| Rucaparib (phosphate) |
| WHI-P97 |
| Cerdulatinib |
| Tenovin-6 |
| ML324 |
| BGP-15 |
| Dihydrocoumarin |
| GSK-690693 |
| CEP-33779 |
| SID 3712249 |
| O-304 |
| I-BRD9 |
| WP1066 |
| EED226 |
| GSK3326595 |
| SGI-1027 |
| AK-7 |
| SR-4370 |
| GSK-J4 |
| UNC0379 |
| MG 149 |
| Metformin (hydrochloride) |
| Mitoxantrone |

| |
|---|
| Santacruzamate A |
| OF-1 |
| Bromosporine |
| Birabresib |
| C-7280948 |
| Dacinostat |
| E7449 |
| TAK-632 |
| Sodium phenylbutyrate |
| Hesperadin |
| Pinometostat |
| IOX1 |
| Alisertib |
| Tucidinostat |
| EPZ011989 |
| NI-57 |
| CPI-169 racemate |
| UPF 1069 |
| Reversine |
| WHI-P154 |
| RGFP966 |
| SB1317 |
| BAY-299 |
| ZM39923 (hydrochloride) |
| Dorsomorphin (dihydrochloride) |
| UNC 669 |
| Baricitinib (phosphate) |
| PFI-1 |
| Resveratrol |
| ZL0420 |
| PLX51107 |
| PF-9363 |
| Bempedoic acid |
| HTH-01-015 |
| Sinapinic acid |
| Tasquinimod |
| EPZ004777 |
| Fimepinostat |
| BG45 |

| |
|---|
| Valproic acid (sodium salt) |
| UF010 |
| Vorinostat |
| 3-TYP |
| Apabetalone |
| GSK2801 |
| Mitoxantrone (dihydrochloride) |
| MS436 |
| Ruxolitinib (phosphate) |
| AICAR |
| M344 |
| Sodium Butyrate |
| Bufexamac |
| BRD73954 |
| Flufenamic acid |
| Momelotinib sulfate |
| Tubastatin A |
| TG101209 |
| Droxinostat |
| HA-100 |
| CPI-455 |
| MN-64 |
| Hinokitiol |
| Belinostat |
| UNC0642 |
| WT-161 |
| BMS-911543 |
| Peficitinib |
| CPI-637 |
| AZD-1480 |
| MI-3 |
| A-966492 |
| 3-Aminobenzamide |
| Enzastaurin |
| Scriptaid |
| UNC-926 |
| Sulforaphane |
| Cambinol |
| AT9283 |

| |
|---|
| BCI-121 |
| Menin-MLL inhibitor MI-2 |
| Fisetin |
| JANEX-1 |
| PHA-680632 |
| AZD5153 (6-Hydroxy-2-naphthoic acid) |
| AMI-1 |
| PF-CBP1 hydrochloride |
| BET bromodomain inhibitor |
| AK-1 |
| AZD1152 |
| Tozasertib |
| PF-06726304 |
| KG-501 |
| Barasertib-HQPA |
| NKL 22 |
| JNJ-7706621 |
| XL228 |
| Selisistat |
| Remetinostat |
| UNC0646 |
| TMP269 |
| UNC0638 |
| Veliparib (dihydrochloride) |
| Fedratinib |
| Tofacitinib (citrate) |
| MM-102 (TFA) |
| TCS7010 |
| CCT 137690 |
| UNC3866 |
| GSK503 |
| WZ4003 |
| Gandotinib |
| BI 2536 |
| PFI-3 |
| γ-Oryzanol |
| UNC0224 |
| BIX-01294 |

| |
|---|
| Givinostat (hydrochloride monohydrate) |
| GSK126 |
| OICR-9429 |
| Tazemetostat |
| 666-15 |
| UNC1999 |
| G007-LK |
| CGP60474 |
| JW 55 |
| I-BET151 |
| Ingenol |
| Molibresib |
| HDAC8-IN-1 |
| CHZ868 |
| Sotrastaurin |
| BI-7273 |
| UNC1215 |
| SNS-314 |
| TMP195 |
| GSK1324726A |
| ITSA-1 |
| C646 |
| Rucaparib (Camsylate) |
| Domatinostat |
| Curcumin |
| (R)-(-)-JQ1 Enantiomer |
| Amodiaquin (dihydrochloride dihydrate) |
| XAV-939 |
| CUDC-101 |
| PF-06409577 |
| Mocetinostat |
| RG2833 |
| PJ34 (hydrochloride) |
| GSK-J1 |
| TAS-301 |
| Solcitinib |
| GSK-5959 |
| Sirtinol |

| |
|---|
| Pracinostat |
| MC1568 |
| Filgotinib |
| Danthron |
| Lomeguatrib |
| AS8351 |
| JIB-04 |
| Anacardic Acid |
| CeMMEC1 |
| Salermide |
| CAY10602 |
| EI1 |
| PJ34 |
| ZLN024 (hydrochloride) |
| Mivebresib |
| SAR-20347 |
| Momelotinib |
| A-366 |
| PCI-34051 |
| BRD4770 |
| AICAR (phosphate) |
| SP2509 |
| Go 6983 |
| BI-9564 |
| SGC707 |
| OSS_128167 |
| GSK2879552 |
| Ricolinostat |
| AZD-2461 |
| LMK-235 |
| BML-210 |
| CPI-203 |
| RG108 |
| GSK591 |
| MS049 |
| HLCL-61 (hydrochloride) |
| Pimelic Diphenylamide 106 |
| WDR5-0103 |
| GSK6853 |

| |
|---|
| Remodelin (hydrobromide) |
| CARM1-IN-1 (hydrochloride) |
| Selisistat R-enantiomer |
| GLPG0634 analog |
| HPOB |
| NVP-TNKS656 |
| EPZ020411 (hydrochloride) |
| JQ-1 (carboxylic acid) |
| YF-2 |
| BET-IN-1 |
| MS417 |
| Niraparib R-enantiomer |
| HDAC-IN-3 |
| GSK4028 |
| DC_517 |
| Baricitinib |
| Upadacitinib |
| DC-05 |
| Sirt2-IN-1 |
| (S)-JQ-35 |
| Y06036 |
| RGB-286638 (free base) |
| Valrubicin |
| EDO-S101 |
| Bisindolylmaleimide I |
| MI-538 |
| CARM1-IN-1 |
| Itacitinib |
| T-3775440 hydrochloride |
| BRD4 degrader AT1 |
| Ilginatinib (maleate) |
| SF2523 |
| CPI-360 |
| MI-463 |
| PKC-IN-1 |
| ABBV-744 |
| (+)-JQ1 PA |
| BRD7-IN-1 |
| CAY10603 |

| |
|---|
| Ruxolitinib (S enantiomer) |
| BETd-260 |
| Corin |
| Tubacin |
| NCGC00244536 |
| MIR96-IN-1 |
| EPZ015666 |
| TH34 |
| KDM5-IN-1 |
| (+)-JQ-1 |
| Tubastatin A (Hydrochloride) |
| GSK 690 (Hydrochloride) |
| Veliparib |
| PF-03814735 |
| ACY-775 |
| Decitabine |
| UNC0321 |
| GNE-049 |
| JAK3-IN-1 |
| IACS-9571 Hydrochloride |
| I-CBP112 |
| VTX-27 |
| MC3482 |
| Danusertib |
| GNE-272 |
| SW-100 |
| MI-503 |
| Abrocitinib |
| EX229 |
| TyK2-IN-2 |
| KDM4D-IN-1 |
| Tyk2-IN-5 |
| TPOP146 |
| MLN8054 |
| MK-8745 |
| DDP-38003 (trihydrochloride) |
| HJB97 |
| GSK 4027 |
| Oxamflatin |

| |
|---|
| LXS196 |
| JNJ-64619178 |
| Zebularine |
| PKC-theta inhibitor |
| Pacritinib |
| Y06137 |
| Sirtuin modulator 1 |
| PF-06700841 (P-Tosylate) |
| BAY1238097 |
| NSC 228155 |
| XY1 |
| SCH-1473759 (hydrochloride) |
| PROTAC Sirt2 Degrader-1 |
| INCB-057643 |
| Talazoparib (8R,9S) |
| KDM5A-IN-1 |
| Histone Acetyltransferase Inhibitor II |
| BAY-598 |
| HDACs/mTOR Inhibitor 1 |
| XL019 |
| Tenovin-1 |
| Olaparib |
| AZ505 (ditrifluoroacetate) |
| GSK9311 |
| SGC-CBP30 |
| SKLB-23bb |
| Midostaurin |
| YLF-466D |
| Trichostatin A |
| Bisindolylmaleimide X (hydrochloride) |
| Niraparib hydrochloride |
| Quisinostat |
| Doxorubicin (hydrochloride) |
| dBET6 |
| Talazoparib tosylate |
| Talazoparib |
| FL-411 |
| NSC 42834 |
| PFI-2 (hydrochloride) |

| |
|---|
| AZ6102 |
| GeA-69 |
| ZXH-3-26 |
| Staurosporine |
| QC6352 |
| GSK-LSD1 Dihydrochloride |
| FM381 |
| MK-5108 |
| BMS-986165 |
| ARV-825 |
| ACY-738 |
| A-485 |
| MZ 1 |
| Delgocitinib |
| MZP-54 |
| WM-8014 |
| BB-Cl-Amidine (hydrochloride) |
| PROTAC BRD9 Degrader-1 |
| 7-Methoxyisoflavone |
| SIRT5 inhibitor 1 |
| Ilginatinib hydrochloride |
| Selisistat S-enantiomer |
| Cucurbitacin I |
| Benzenebutyric acid |
| MK8722 |
| JQEZ5 |
| Delcasertib |
| ORY-1001(trans) |
| GSK484 (hydrochloride) |
| PF-06651600 |
| DCP-LA |
| GSK-1070916 |
| RK-287107 |
| GSK2807 Trifluoroacetate |
| ACY-957 |
| MK-3903 |
| MZP-55 |
| Pyridone 6 |
| (3S,4S)-Tofacitinib |

| |
|---|
| Glucose-conjugated MGMT inhibitor |
| PROTAC BET Degrader-1 |
| Guadecitabine sodium |
| KW-2449 |
| Ginkgolide C |
| Itacitinib adipate |
| Gusacitinib |
| dTRIM24 |
| dBET57 |
| AMG 900 |
| EZM 2302 |
| ENMD-2076 |
| INCB054329 |
| Ilginatinib |
| D-erythro-Sphingosine |
| CPI-169 |
| CCT241736 |
| Phorbol 12,13-dibutyrate |
| NCGC00247743 |
| UNC 0631 |
| JAK/HDAC-IN-1 |
| Seclidemstat |
| Daphnetin |
| Ruxolitinib |
| Iniparib |
| LFM-A13 |
| Entinostat |
| Nexturastat A |
| Tofacitinib |
| Valproic acid |
| EPZ005687 |
| CF53 |
| ME0328 |
| Nicotinamide riboside (chloride) |
| UBCS039 |
| CBP-IN-1 |
| MS023 (dihydrochloride) |
| ZM-447439 |
| BETd-246 |

| |
|---|
| 5-Azacytidine |
| AZ960 |
| GNE-781 |
| FLLL32 |
| GSK467 |
| SGC-iMLLT |
| SGC2085 |
| LY3295668 |
| Cl-amidine (hydrochloride) |
| GSK343 |
| BRD 4354 (ditrifluoroacetate) |
| CXD101 |
| EPZ031686 |
| Citarinostat |
| A1874 |
| ARV-771 |
| (-)-Indolactam V |
| Molibresib besylate |
| (rac)-BAY1238097 |
| Tacedinaline |
| Niraparib |
| (R)-BAY1238097 |
| Thiomyristoyl |
| 5-Methyl-2'-deoxycytidine |

**Supplementary Table S1:** List of epigenetic modifiers used for drug screen. Compounds purchased from Med Chem Express.

| sgRNA | Target Sequence |
|---|---|
| KAT6A clone 1 | ACAGCTCAAGAAAAGCCCTG |
| KAT6A clone 2 | TTTGGCATACGGGTGAGTGG |
| KAT6B clone 1 | ATATTCCTGTGGGTAAGGCG |
| KAT6B clone 2 | TCTTCATCTGAGTTGTCAAA |

**Supplementary Table S2.** Oligo sequences: Sequence information for oligos used as sgRNA for CRISPR-mediated gene disruption.

| Antibody | Assay | Manufacturer | Catalog # | Lot# |
|---|---|---|---|---|
| H3K23ac | ChIP-seq | Abcam | 177275 | GR3358483-7 |
| H3K27ac | ChIP-seq | Abcam | 177178 | 1041853-1 |
| H3K27me3 | Cut&Run | Abcam | 192985 | 1068234-1 |
| KAT6A | Cut&Run/WB | Millipore | 065052 | 10208 |
| KAT6B | Cut&Run/WB | Millipore | 006104 | 000023886 |
| IgG | Cut&Run | Santa Cruz | Sc-2028 | H2613 |
| MYCN | WB | Cell signaling | 9405S | 2 |
| PHOX2B | WB | Santa Cruz | Sc-376993 | E3012 |
| GATA3 | WB | Cell signaling | 5852 | 5 |
| actin | WB | Cell signaling | 4070 | 18 |
| Caspase 3 | WB | Cell signaling | 9661S | 47 |
| PARP | WB | Cell signaling | 9542S | 15 |
| Total H3 | WB | Active Motif | 39763 | 0301 |
| H3K23ac | WB | Active Motif | 39131 | 34519001 |
| H3K9ac | WB | Cell signaling | 9649T | 13 |
| H3K27ac | WB | Cell signaling | 39133 | 31814008 |
| Ki67 | IHC | Biocare | CRM325B | 101420A-2 |
| GD2 | IHC | BD Pharmingen | 554272 | 3276204 |
| GD2 | Flow cytometry | Santa Cruz | Sc-53831 | K0922 |
| Secondary antibody | Flow cytometry | Invitrogen | A27040 | 2649807 |

**Supplementary Table S3.** Antibody information: The antibodies used for ChIP-seq, CUT&Run, western blot (WB), Immunohistochemistry (IHC) and flow cytometry are listed along with the manufacturer, catalog number and specific lot number.

| Cell line | Assay | Treatment | Target | GEO Accession # |
|---|---|---|---|---|
| BE2C | ChIP-seq | none | H3K23ac | pending |
| BE2C | ChIP-seq | none | input | pending |
| BE2C | ChIP-seq | none | H3K27ac | pending |
| BE2C | ChIP-seq | none | input | pending |
| BE2C | ATAC- seq | none | N/A | GSM2486171 |
| BE2C | CUT& RUN | DMSO | H3K27me3 | pending |
| BE2C | CUT& RUN | Isotretinoin | H3K27me3 | pending |
| BE2C | CUT& RUN | PF-9363 | H3K27me3 | pending |
| BE2C | CUT& RUN | PF-9363+Isotretinoin | H3K27me3 | pending |
| BE2C | CUT& RUN | none | KAT6A | pending |
| BE2C | CUT& RUN | none | KAT6B | pending |
| BE2C sgNT1 | RNA-seq | none | N/A | pending |
| BE2C sgKAT6A | RNA-seq | none | N/A | pending |
| BE2C sgKAT6B | RNA-seq | none | N/A | pending |
| BE2C, repl 1 | RNA-seq | DMSO | N/A | pending |
| BE2C, repl 2 | RNA-seq | DMSO | N/A | pending |
| BE2C, repl 3 | RNA-seq | DMSO | N/A | pending |
| BE2C, repl 1 | RNA-seq | Isotretinoin | N/A | pending |
| BE2C, repl 2 | RNA-seq | Isotretinoin | N/A | pending |
| BE2C, repl 3 | RNA-seq | Isotretinoin | N/A | pending |
| BE2C, repl 1 | RNA-seq | PF-9363 | N/A | pending |
| BE2C, repl 2 | RNA-seq | PF-9363 | N/A | pending |
| BE2C, repl 3 | RNA-seq | PF-9363 | N/A | pending |
| BE2C, repl 1 | RNA-seq | PF-9363+ Isotretinoin | N/A | pending |
| BE2C, repl 2 | RNA-seq | PF-9363+ Isotretinoin | N/A | pending |
| BE2C, repl 3 | RNA-seq | PF-9363+ Isotretinoin | N/A | pending |
| BE2C | scRNA-seq | DMSO | N/A | pending |
| BE2C | scRNA-seq | Isotretinoin | N/A | pending |
| BE2C | scRNA-seq | PF-9363 | N/A | pending |
| BE2C | scRNA-seq | PF-9363+ Isotretinoin | N/A | pending |

**Supplementary Table S4.** NCBI GEO accession numbers; raw and processed data files are deposited to the NCBI GEO server under super-series GSEXXXX.
